# A cellular and molecular atlas reveals the basis of chytrid development

**DOI:** 10.1101/2021.09.06.459148

**Authors:** Davis Laundon, Nathan Chrismas, Kimberley Bird, Seth Thomas, Thomas Mock, Michael Cunliffe

**Author notes:** Correspondence: Michael Cunliffe Marine Biological Association, The Laboratory, Citadel Hill, Plymouth, PL1 2PB, UK. E.

## Abstract

The chytrids (phylum Chytridiomycota) are a major early-diverging fungal lineage of ecological and evolutionary importance. Despite their importance, many fundamental aspects of chytrid developmental and cell biology remain poorly understood. To address these knowledge gaps, we combined quantitative volume electron microscopy and comparative transcriptome profiling to create an ‘atlas’ of the cellular and molecular basis of the chytrid life cycle, using the model chytrid *Rhizoclosmatium globosum*. From our developmental atlas, we show that zoospores exhibit a specialised biological repertoire dominated by inactive ribosome aggregates, and that lipid processing is complex and dynamic throughout the cell cycle. We demonstrate that the chytrid apophysis is a distinct subcellular structure characterised by high intracellular trafficking, providing evidence for division of labour in the chytrid cell plan, and show that zoosporogenesis includes ‘animal like’ amoeboid cell morphologies resulting from endocytotic cargo transport from the interstitial maternal cytoplasm. Taken together, our results reveal insights into chytrid developmental biology and provide a basis for future investigations into early-diverging fungal cell biology.

## INTRODUCTION

The chytrids (phylum Chytridiomycota) are an early-diverging and predominantly unicellular fungal lineage of ecological importance. For example, some chytrid species are the causative agents of the global amphibian panzootic (Fisher and Garner, 2020) and virulent crop pests (van de Vossenberg et al., 2019), whilst others are algal parasites and saprotrophs in marine and freshwater ecosystems (Frenken et al., 2017; Grossart et al., 2019; Klawonn et al., 2021). Chytrid zoospores contain large amounts of intracellular storage lipids that are consumed by grazing zooplankton, making them responsible for a significant form of trophic upgrading in aquatic ecosystems (Kagami et al., 2017, 2014, 2007; Rasconi et al., 2020). Chytrids are also important from an evolutionary perspective as they retain cellular traits from the last common ancestor of branching fungi (Fig. 1A) that are now absent in hyphal fungi (Berbee et al., 2017; Nagy et al., 2018), as well as traits from the common ancestor of animals and fungi in the Opisthokonta (Medina et al., 2016; Prostak et al., 2021). This makes chytrids powerful models to explore the origin and evolution of innovations in fungal cell biology and the wider eukaryotic tree of life. To help fully appreciate chytrids in terms of their ecological and evolutionary contexts, it is necessary to resolve their core cell biology (Laundon and Cunliffe, 2021).

**Figure 1.**
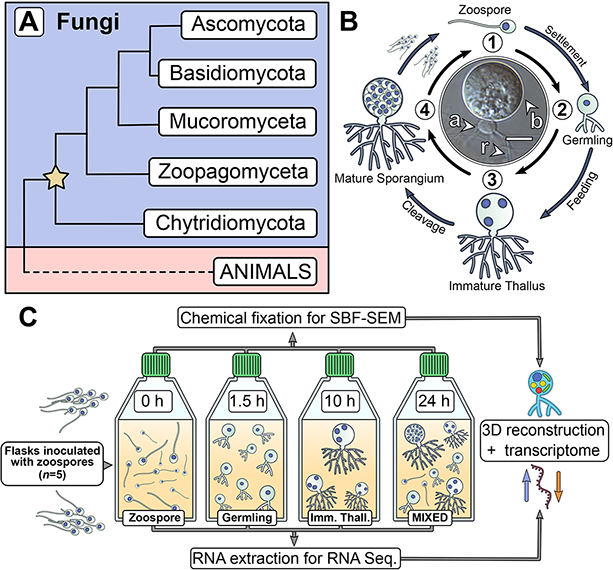
Chytrids are an early-diverging fungal phylum with a dimorphic life cycle. (A) Chytrids (phylum Chytridiomycota) are an early-diverging fungal lineage, many members of which exhibit cellular characteristics retained from the last common ancestor of branching (rhizoidal and hyphal) fungi (star). Simplified phylogenetic tree from (Laundon and Cunliffe, 2021; Tedersoo et al., 2018). (B) *Rhizoclosmatium globosum* exhibits an archetypal chytrid life cycle and cell plan delineated here into four discrete major stages. Labelled is the apophysis (a), cell body (b), and rhizoids (r). Scale bar = 10 µm. (C) Diagrammatic workflow of the experimental setup used in this study for comparative cellular Serial Block Face Scanning Electron Microscopy (SBF-SEM) and molecular (transcriptome) analysis.

Central to chytrid cell biology is their distinctive dimorphic life cycle, consisting of a motile free-swimming uniflagellate zoospore that transforms into a sessile walled thallus with anucleate attaching and feeding rhizoids (Fig. 1B) (Laundon et al., 2020; Medina et al., 2019). The cell body component of the thallus develops into the zoosporangium from which the next generation of zoospores are produced (Fig. 1B). Any biological life cycle inherently represents a temporal progression, yet the chytrid life cycle can be categorised into four distinctive contiguous life stages (Berger et al., 2005). The first stage is the motile ‘zoospore’ which lacks a cell wall, does not feed, and colonises substrates or hosts. The second stage is the sessile ‘germling’ which develops immediately after zoospore settlement following flagellar retraction (or sometimes detachment), cell wall production (encystment), and initiation of rhizoid growth from an initial germ tube. The third stage is the vegetative ‘immature thallus’ which is associated with the highest levels of rhizoid development and overall cellular growth (Laundon et al., 2020). The cell plan of the immature thallus can be divided into three parts (Fig. 1B): the cell body which is ultimately destined for reproduction (zoosporogenesis), the rhizoid for attachment and feeding, and (in some chytrid species) a bulbous swelling between the cell body and rhizoid termed the ‘apophysis’, the function of which is currently poorly understood (Laundon and Cunliffe, 2021). The final life stage is the reproductive ‘mature zoosporangium’, which appears once the immature thallus has reached maximum cell size and the cell body cytoplasm is cleaved into the next generation of zoospores (Fig. 1B).

Representative model strains have an important role to play in understanding the biology of chytrids (Laundon and Cunliffe, 2021). *Rhizoclosmatium globosum* is a widespread aquatic saprotroph and the strain *R. globosum* JEL800 has emerged as a promising model organism in laboratory investigations (Laundon et al., 2020; Roberts et al., 2020; Venard et al., 2020) due to an available genome (Mondo et al., 2017), easy axenic culture, and amenability to live-cell fluorescent microscopy (Laundon et al., 2020). *R. globosum* JEL800 exhibits an archetypal chytrid cell plan (Fig. 1B) and rapid life cycle, making it a useful model system to interrogate cellular development (Laundon et al., 2020; Laundon and Cunliffe, 2021).

The chytrid cell cycle has so far been characterised with largely descriptive approaches (e.g Berger et al., 2005) with only a few quantitative studies focusing on specific processes, such as rhizoid morphogenesis (Dee et al., 2019; Laundon et al., 2020) and actin formation (Medina et al., 2020; Prostak et al., 2021). These important studies provide a foundation on which to develop a quantitative approach to understand the biology of the chytrid cell plan and the drivers of the transitions between the life stages. To investigate the cellular and molecular underpinnings of the chytrid life cycle and associated cell biology, we studied the four major life stages of the *R. globosum* cell cycle by combining quantitative volume electron microscopy and transcriptomics (Fig. 1C), with the addition of supplementary targeted live-cell fluorescent microscopy and lipid analysis. The aim of our approach was to quantify the cellular traits that define the major chytrid life stages and identify the biological processes that take place during the developmental transitions between them. As such, we have created a developmental ‘atlas’ with *R. globosum* for the archetypal chytrid lifecycle, which in turn generated specific avenues for targeted investigation of important biological processes, namely lipid biology, apophysis function, and zoosporogenesis.

By culturing *R. globosum* and sampling the populations at different stages through their temporal development (0 h zoospore, 1.5 h germling, 10 h immature thallus, and a 24 h mixed population including mature zoosporangia) (Fig. 1C), we examined chytrid populations with both 3D reconstructions by Serial Block Face Scanning Electron Microscopy (SBF-SEM) and mRNA sequencing. We used single-cell SBF-SEM reconstructions (*n* = 5) (Suppl. Fig. 1) to quantify the cellular structures at each life stage and population-level transcriptomic analysis of significant KEGG pathway categories (*n* = 3, Differentially Expressed Genes (DEGs)) to identify the major biological differences between the life stages through temporal development (Fig. 1C). As these stages represent key time points in the progression of the linear temporal chytrid life cycle, pairwise comparison of transcriptomes from contiguous life stages achieved an account of the major putative biological transitions (e.g. germling vs. zoospore, immature thallus vs. germling) (Fig. 2C; Suppl. Figs 2-5). Our findings provide insights into chytrid developmental processes and serve as a resource from which to resolve the biology of this ecologically and evolutionary important fungal lineage.

**Figure 2.**
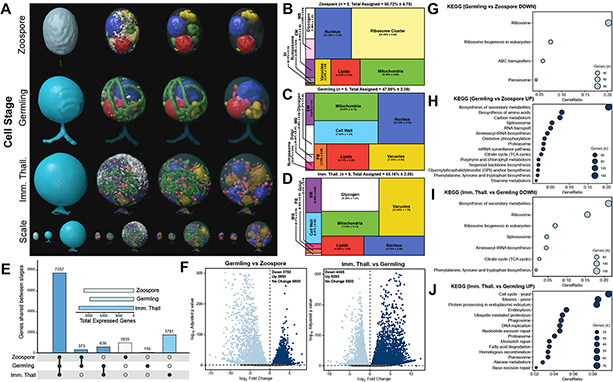
Serial Block Face Scanning Electron Microscopy (SBF-SEM) reconstructions and transcriptome analysis provided an atlas of the *Rhizoclosmatium globosum* life cycle. (A) Representative SBF-SEM reconstructions of the first three life stages of the *R. globosum* lifecycle. Bottom row shows the stages to scale. Organelle colours as in (B-D) and conserved throughout. (B-D) Volumetric composition of assigned organelles in SBF-SEM reconstructions (*n* = 5) of zoospores (B), germlings (C), and immature thalli (D). EM = endomembrane, MB = microbodies, PB = peripheral bodies, SI = striated inclusion. (E) Shared and unique gene expression counts between life stages. Inset shows total expressed genes per life stage. (F) Pairwise comparison of differentially expressed genes (DEGs) between germlings and zoospores, and immature thalli and germlings. (G-J) Pairwise comparison of significant (*p* < 0.05) differentially expressed KEGG categories between germlings and zoospores (G-H), and immature thalli and germlings (I-J).

## RESULTS

### A cellular and molecular atlas of *Rhizoclosmatium globosum*

The orientation, subcellular localisation, and morphology of the cellular ultrastructure determined with SBF-SEM of the *R. globosum* zoospore, germling, and immature thallus life stages are shown in (Fig. 2A; Movie 1; Suppl. Mov. 1-21; Suppl. File 1), with the volumetric transition from zoospore (20.7±1.7 µm^3^) to germling (33.0±2.0 µm^3^) to immature thallus (1,116.3±206.2 µm^3^) exceeding an order of magnitude. These ultrastructural differences in the cell patterns at each life stage (Fig. 2B-D) are complemented with differential gene expression analysis focusing on characterising the transitions between life stages (Fig. 2E-J). Full statistical details of cell volumetric and molecular comparisons are provided in (Suppl. Fig. 5-7; Suppl Tables 1-5; Suppl. File 2). As the mature zoosporangium samples were taken from a mixed population of cell stages (Fig. 1C), they were conservatively excluded from comparison with the first three stages and will be treated separately in this analysis.

The zoospore cell body is a prolate spheroid with an apical flagellum and is volumetrically dominated by a structurally distinct ribosome cluster (20.5±2.8 %) in the cell interior which was not detected in the other life stages (Fig. 2A-B; Suppl. Fig. 2; Suppl. Mov. 1-5). The loss of the ribosome cluster in the germling from the zoospore stage matched a downregulation of ribosome and ribosome biogenesis KEGG categories in the germling relative to the zoospore (Fig. 2G; Suppl. File 2). There was no observed significant molecular signature associated with elevated protein synthesis in zoospores, suggesting that ribosome presence and aggregation, but not activity, govern chytrid zoospores. Only two other KEGG categories were downregulated in germlings relative to zoospores. These were linked to peroxisomes and ATP-binding cassette (ABC) transporters (Fig. 2G), both of which are associated with lipid metabolism (discussed further below).

Following encystment, the germling stage marks the origin of a complete cell wall, Golgi apparatuses (3 out of 5 replicates), and peripheral bodies i.e. vesicular structures bound to the cell periphery putatively associated with cell wall deposition (Fig. 2A&C; Suppl. Fig. 2; Suppl. Mov. 6-10), as well as the beginning of rhizoid growth from an apical germ tube. Transcriptome analysis indicated that the germling exhibits a greater range of active processes compared to the zoospore, with upregulation of primary and secondary metabolism (e.g. amino acid and secondary metabolite biosynthesis), feeding and energy release (e.g. carbon metabolism and Tricarboxylic Acid Cycle), and transcription and translation (e.g. spliceosome and aminoacyl-tRNA biosynthesis) KEGG categories (Fig. 2H). A similar pattern is shown when comparing KEGG categories downregulated in immature thalli relative to germlings (Fig. 2I). In the germling stage, we also show upregulation of genes associated with proteasome activity (Fig. 2H). Taken together, these data show that the transition from zoospore to germling is characterised by the apparent activation of diverse biological processes including central metabolic pathways, cellular anabolism, and feeding.

Compared to the germling, immature thalli devoted a smaller volumetric proportion to the cell wall (IT 2.4±0.3 % vs G 7.6±1.2 %, *p*<0.01) and peripheral bodies (IT 0.3±0.1 % vs G 1.7±0.3 %, *p*<0.01) (Fig. 2A&D; Suppl. Fig. 5; Suppl. Mov. 11-15). Similarly, nuclei (IT 4.8±2.5 % vs G 12.2±0.5 %, *p*<0.01) and mitochondria (IT 7.0±0.1 % vs G 9.1±0.7%, *p*<0.001) occupied a smaller volumetric proportion (Fig. 2A&D; Suppl. Fig. 5). Conversely, immature thalli displayed larger glycogen stores (IT 9.4±2.0 % vs G 1.3±0.4%, *p*<0.01) and vacuole fractions (IT 13.0±1.8 % vs G 7.6±0.9 %, *p*<0.001) than germlings (Fig. 2A&D; Suppl. Fig. 5). Quantification of the increased vacuolisation of immature thalli in the SBF-SEM reconstructions was correlated with upregulation of related KEGG categories including endocytosis and phagosomes relative to germlings (Fig. 2J). Within these categories are genes related to microtubules and actin, including actin-related proteins-2/3 (ARP2/3), indicating that the immature thalli are associated with higher cytoskeletal activity compared to germlings. Some immature thallus replicates were multinucleate (1.8±1.3 nuclei per cell), indicating the onset of nuclear division (Fig. 2A), which matched the upregulation of cell cycle and DNA replication KEGG categories relative to germlings. The apophysis (12.2±6.0 µm^3^) was observed at the immature thallus stage (discussed further below). Overall, these data show that the biological shift from germling to immature thallus is characterised by a move from initiating general metabolic activity to intracellular trafficking and the start of zoosporogenesis.

As anticipated, the SBF-SEM reconstructions showed that the zoospore is wall-less unlike the germling and immature thallus stages (Fig. 2). Single cell fluorescent-labelling of chitin (the primary wall component) however showed that precursory material is produced by zoospores at the apical pole near the flagellum base (Fig 3A) suggesting that cell wall production is initiated to some extent during the free-swimming zoospore stage of the *R. globosum* cell cycle. In a previous study (Laundon et al., 2020), we identified twenty-eight candidate genes for glycosyltransferase (GT2) domain-containing proteins putatively involved in chitin synthesis in *R. globosum* and searched for their individual regulation in the transcriptome data. There was no clear pattern of differential regulation of these genes between the life stages overall, however five putative chitin synthase genes were upregulated during the zoospore stage (Suppl. File 2). Nine genes were only found upregulated in the thallus relative to the germling, six of which had >5-fold change increase in abundance. Six genes were not recovered in any the transcriptomes. Interestingly, a putative β-1,6-glucan synthase gene (ORY39038) identified in (Laundon et al., 2020) as having a possible role in wall formation in *R. globosum* was downregulated in germlings relative to wall-less zoospores. Together, this suggests that cell wall formation is a dynamic process throughout the chytrid life cycle, with alternative synthesis enzymes employed at different stages.

**Figure 3.**
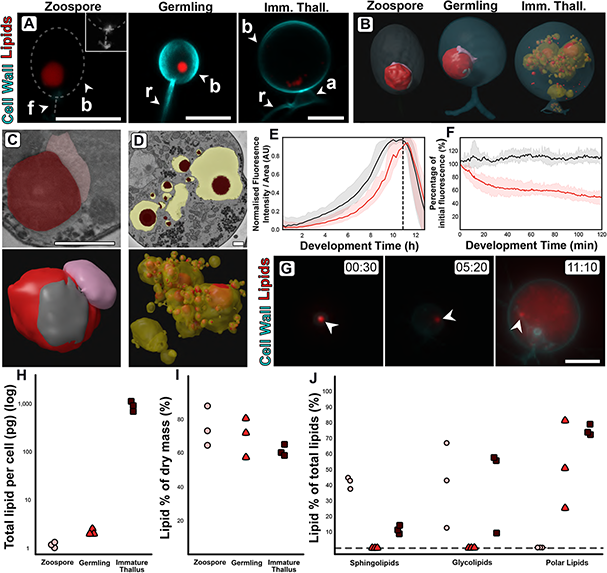
Changes in lipid and lipid-associated cell structures occur with transitions between *R. globosum* life stages. (A) Fluorescent labelling of *R. globosum* shows distinct shifts in lipid structures across the chytrid life cycle and cell wall. Dashed line demarks cell boundary where not labelled in the zoospore. Zoospore inset shows precursory cell wall material at the flagellar base contrast-brightness adjusted for visualisation. Apophysis (a), cell body (b), flagellum (f), rhizoid (r). Scale bars = 5 µm. (B) Representative SBF-SEM reconstructions of lipid globules and lipid-associated structures across chytrid life stages. (C-D) Representative single false-coloured SBF-SEM slices (top) and SBF-SEM reconstructions (bottom) of the lipid-rumposome-microbody (LRM) complex from zoospores (also seen in germlings) (C) and intravacuolar lipid globules (D) from immature thalli. Scale bars = 1 µm. (E) Live-cell imaging (*n* = 5) of *R. globosum* population-level Nile red-stained lipid dynamics. Red = mean lipid fluorescence (± min/max), black = mean total cell area (± min/max), dashed line = mean sporulation time of population. (F) Immediately following zoospore settlement, the population-level (*n* = 5) lipid fluorescence (red) decreases relative to fixed photobleaching control populations (black). (G) Live-cell imaging revealed differential lipid dynamics across the chytrid life cycle. Note that the original zoospore lipid globule (arrowhead) remains intact up to the point of lipid anabolism in the immature thallus. Timestamp = HH:MM. Scale bar = 10 µm. (H-J) Lipid analysis shows shifts in lipid composition of the chytrid lifecycle. Lipid quantities as total mass per cell (H) and as a percentage of total dry mass (I) between chytrid life stages. Changes in lipid fractions were found between chytrid life stages (J). Dashed line = below analytical detection.

### Changes in subcellular lipid-associated structures are linked with variation in lipid composition

Fluorescent labelling and SBF-SEM reconstructions showed that zoospores and germlings possess a single lipid globule (Z 0.9±0.6 and G 1.9±1.1 µm^3^) whereas immature thalli have multiple (68.8±55.2) but smaller (0.5±0.8 µm^3^) globules scattered throughout the cell body (Fig. 3A-B; Movie 2). The lipid globule (red) in the zoospore and germling stages was associated with an apically oriented structure called the rumposome (grey), which is a chytrid-specific organelle putatively associated with cell signalling (Powell, 1983), and a basally oriented microbody (pink) that likely functions as a lipid-processing peroxisome (Powell, 1976) (Fig. 3B-C; Movie 2). Together these structures form the lipid-rumposome-microbody (LRM) complex. The rumposome was larger in zoospores than in germlings (Z 0.3±0.0 % vs G 0.1±0.1 %, *p*<0.001) (Suppl. Fig. 5), indicating increased activity in zoospores. In immature thalli, LRM complexes were not detected. Unlike in zoospores and germlings, the bulk of the lipids in the immature thalli were intravacuolar (89.8±8.5 % total lipids) (Fig. 3D). There was no proportional volumetric difference in lipid fractions determined with SBF-SEM reconstructions between the three life stages (Z 4.3±2.6 % vs G 5.7±3.7 % vs IT 4.0±1.6 %, *p*>0.05) (Fig. 2B-D; Suppl. Fig. 5).

Live-population imaging of Nile Red-labelled storage lipids showed that initially (0-2 h) the chytrid life cycle was characterised by a decrease (−49.7±9.8 %) in lipid fluorescence suggesting that neutral storage lipid catabolism was taking place, before fluorescence increased suggesting that lipid anabolism was occurring up to zoospore release (Fig. 3E-F). The initial lipid fluorescence decreased even in the presence of a carbon replete growth medium in line with the non-feeding habit of zoospores (Fig. 3F). Live-single cell imaging revealed a similar response as shown at the population level, and additionally showed that the zoospore lipid globule remains intact and detectable until at least the point of visible lipid anabolism in the developing cell when the globule becomes undisguisable from the new lipids (Fig. 3G, Movie 3; Suppl. Mov. 22).

Extraction and quantification of lipids from cells harvested at the major life stages showed shifts in lipid profiles. Individual zoospores possessed 1.2±0.1 pg, germlings 2.2±0.3 pg, and immature thalli 904.5±201.0 pg of lipid per cell (Fig. 3H), however lipid composition as a percentage of dry mass (Z 74.8±11.8 % vs G 69.5.0±11.5 % vs IT 61.0±3.3 %) was similar across the life stages (Fig. 3I). Sphingolipids were present in both zoospores and immature thalli (Z 41.6±3.6 % and IT 11.5±2.6 %), but below detection in germlings (Fig. 3J). Likewise, glycolipids were present in both zoospores (40.7±27.1%) and immature thalli (40.6±27.3 %), but below detection in germlings. Conversely, polar lipids were below detection in zoospores yet present in germlings (51.7±27.5 %) and immature thalli (74.0±3.4 %).

The differences in lipid composition between the zoospore and germling stages (Fig. 3J) correlated with higher expression of genes in KEGG categories associated with peroxisome activity and ABC-transporters in zoospores compared to germlings (Fig. 2G). Most of the genes identified under the peroxisome category are involved in lipid oxidation and acyl-CoA metabolism, and therefore likely involved in the catabolic processing of the lipid globule (Suppl. File 2). Lipid reductases were also detected, which have previously been identified with phospholipid anabolism (Lodhi and Semenkovich, 2014) and, together with the increase in endomembrane between germlings and zoospores determined with SBF-SEM (Figure 2B-C), point to increased phospholipid synthesis for membrane production. ABC-transporters are also involved in lipid transport into peroxisomes from lipid stores (Tarling et al., 2013). Together, these results suggest that glycolipid, and potentially sphingolipid, catabolism likely form part of storage lipid utilisation from the globule via the peroxisome, and polar lipid anabolism for endomembrane production are biological characteristics of the transition from zoospore to germling.

We also observed the upregulation of genes associated with fatty acid degradation and peroxisomes in the immature thallus stage compared to the germling stage (Fig. 2J) coinciding with new lipids being produced (Fig. 3). The genes were associated with similar acyl-CoA pathways as the zoospore peroxisome category, in addition to alcohol and aldehyde dehydrogenation (Suppl. File 2). Interestingly, although the peroxisome category was also upregulated in immature thalli, the associated genes were not identical to those in zoospores. Many similar acyl-CoA metabolic signatures were shared (16 genes), but with the addition of alcohol and isocitrate dehydrogenation and superoxide dismutase activity. This suggests that in immature thalli lipid production is driven by an interplay of fatty acid degradation and lipid anabolism, illustrating that some aspects of lipid catabolism and conversion in zoospores are bidirectionally repurposed for anabolism in immature thalli.

### The apophysis is a compartmentalised junction for intracellular trafficking between the rhizoids and cell body

The apophysis is ubiquitous across the Chytridiomycota (James et al., 2006), but the function of the structure is poorly understood (Laundon and Cunliffe, 2021). Here we show that the apophysis exhibits high endomembrane density and active intracellular trafficking between the feeding rhizoids and cell body (Fig. 4). Live-population imaging of FM1-43 labelled endomembrane in *R. globosum* cells (excluding apophysis and rhizoids) showed stability in fluorescence at the beginning of the life cycle (0-2 h), before a constant increase to the point of zoospore release (Fig. 4A). Matching this, SBF-SEM reconstruction revealed that immature thalli devoted a larger proportion of cell body volume to endomembrane than zoospores and germlings (Z 2.3±1.5 % vs G 7.6±0.9 % vs IT 13.0±1.8 %, *p*<0.001), as well as vacuoles (Fig. 2B-D, Fig. 4B; Suppl. Fig. 5). The associated upregulation of KEGG categories such as protein processing in the endoplasmic reticulum (ER) and ubiquitin mediated proteolysis (Fig. 2J) with the transition from the germling to the immature thallus stage suggests that this structural endomembrane is at least in part ER and coupled with protein turnover.

**Figure 4.**
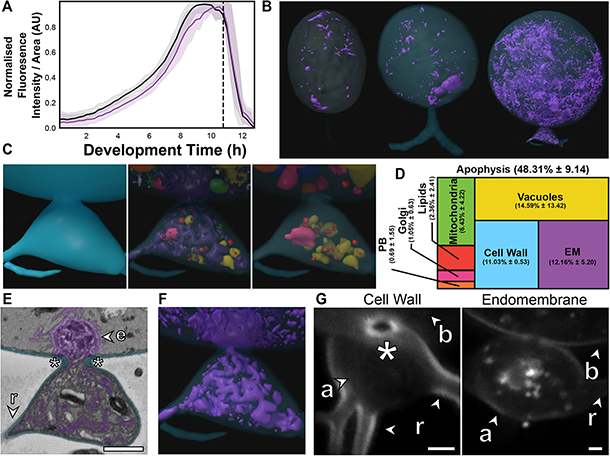
The apophysis is a distinct subcellular structure characterised by increased endomembrane trafficking. (A) Live-cell imaging (*n* = 5) of *R. globosum* population-level FM1-43-stained endomembrane dynamics. Purple = mean endomembrane fluorescence (± min/max), black = mean total cell area (± min/max), dashed line = mean sporulation time of population. (B-C) Representative SBF-SEM reconstructions of endomembrane across chytrid life stages (B) and the apophysis from immature thalli (C). Volumetric composition of SBF-SEM reconstructions (*n* = 5) of immature thallus apophyses (D). Representative single false-coloured SBF-SEM slice (E) and reconstruction (F) of the endomembrane and thickened cell wall (asterisk) at the apophysis-cell body junction. Fluorescent labelling of the chitin rich wall around the apophysis-cell body connecting pore and associated endomembrane structures (G). Labels as in Fig. 3A. All scale bars = 1 µm.

The immature thalli SBF-SEM reconstructions showed that apophyses displayed even greater structural endomembrane than their corresponding cell bodies (apophysis 12.2±5.2 % vs cell body 2.7±0.6 %, *p*<0.01) (Fig. 4C-D; Suppl. Fig. 6; Movie 4). Apophyses also had comparatively more cell wall than the larger cell bodies (A 11.0±0.5 % vs CB 2.4±0.3 %, *p*<0.01) (Fig. 4D; Suppl. Fig. 6). In *R. globosum* the cytoplasm between the apophysis and the cell body is connected via an annular pore (0.40±0.07 µm in diameter) in a distinctive chitin-rich pseudo-septum (Fig. 4 E&G), causing spatial division within the immature thallus cell plan. Live single cell imaging showed dynamic endomembrane activity in the apophysis linking the intracellular traffic between the rhizoid system and apical base of the cell body via the pore (Movie 5; Suppl. Mov. 23). Taken together, we propose that a function of the apophysis is to act as a cellular junction that regulates intracellular traffic and channels material from feeding rhizoids through the pseudo-septal pore to the cell body dedicated for reproduction.

### Developing zoospores in the zoosporangium display an amoeboid morphology resulting from endocytotic trafficking

Understanding zoosporogenesis, including how the thallus differentiates into the next generation of zoospores, is integral to closing the chytrid life cycle. We were unable to achieve a synchronised population of mature zoosporangia, however imaging and sequencing of mixed populations (∼4% cells at the mature zoosporangia stage, Suppl. Fig. 8) still allowed structural characterisation of this life stage, including the SBF-SEM reconstruction of an entire mature zoosporangium containing 82-developing zoospores (Fig. 5; Suppl. Fig. 2; Movie 6; Suppl. Mov. 16). Mature zoosporangia are characterised by internal membrane cleavage (Fig. 5A) where coenocytic immature cytoplasm and organelles are allocated into nascent zoospores. The volume of the SBF-SEM reconstructed mature zoosporangium was 3,651.5 µm^3^, showing that a single chytrid cell volumetrically increases by more than two orders of magnitude over its entire life cycle. The developing zoospores were flagellate, with the flagellum coiled round the cell body in two neat and complete rotations (Fig. 5B). Zoospores are held within the cell wall of the zoosporangium during zoosporogenesis (Fig. 5B), before exiting through the basally oriented discharge pore (an aperture in the cell wall) when developed. During development, the pore is obstructed by a fibrillar discharge plug (49.7 µm^3^ in volume) (Fig. 5C).

**Figure 5.**
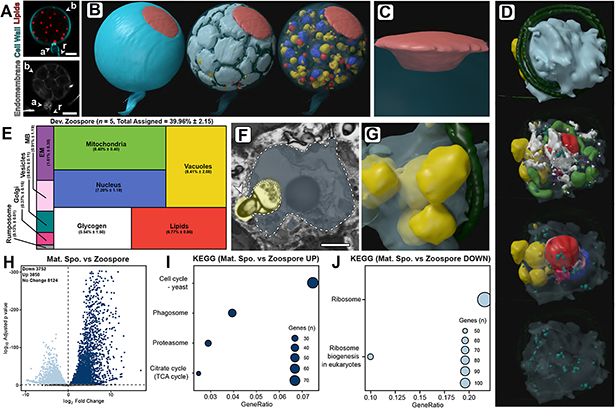
Developing zoospores in the zoosporangium have amoeboid morphology with endocytotic activity. (A) Fluorescent labelling of lipids, cell wall, and endomembrane in an *R. globosum* mature zoosporangium. Scale bar = 5 µm. (B-C) SBF-SEM reconstructions of an 82-zoospore containing mature zoosporangium (B) highlighting the discharge plug, shown in coral (C). (D) Representative SBF-SEM reconstructions of a developing zoospore. Organelle colours as in Fig. 5E. (E) Volumetric composition of SBF-SEM reconstructions of developing zoospores (*n* = 5). (F-G) Representative single false-coloured SBF-SEM slice (F) and reconstruction (G) of the endocytotic vacuoles in developing zoospores. Dashed line delineates the zoospore cell boundary in (F). Scale bar = 1 µm. (H) Pairwise comparison of differentially expressed genes (DEGs) between mature zoosporangia and the free-swimming zoospore life stage. (I-J) Pairwise comparison of significant differentially expressed KEGG categories between mature zoosporangia and the free-swimming zoospore life stage.

The single entire zoosporangium reconstruction (Fig. 5B) allowed the visualisation of developing zoospores in context, but to understand the detailed structural basis of this process it was necessary to reconstruct individual zoospore cells in the zoosporangium at higher resolution for comparison with free-swimming zoospores (Fig. 5D-G; Suppl. Fig. 7; Suppl. Mov. 17-21). This was coupled with comparison of transcriptomes from the mature zoosporangia (taken from the mixed populations) with transcriptomes from the free-swimming zoospores. Relative to mature free-swimming zoospores, developing zoospores in the zoosporangium displayed an amoeboid morphology and had greater intracellular trafficking, characterised by a larger volumetric proportion of endomembrane (DZ 1.7±0.3 % vs MZ 0.9±0.4 %, *p*<0.05), vacuoles, (DZ 8.4±2.1 % vs MZ 2.3±1.5 %, *p*<0.001) and the presence of Golgi apparatuses and a vesicle class not observed in mature free-swimming zoospores (Fig. 5D-E; Suppl. Fig. 7). Developing zoospores in the zoosporangium also exhibited larger glycogen stores (DV 5.5±1.5 % vs MV 1.6±1.2 %, *p*<0.01), indicating that glycogen utilisation occurs between the two stages, and a smaller rumposome (DV 0.1±0.0 % vs MZ 1.3±0.0 %, *p*<0.001) (Fig 5D-E) than their mature free-swimming counterparts.

The amoeboid morphology of the developing zoospores was in part a result of endocytotic engulfment activity, where vacuoles extended from within the zoospore cell interior to the surrounding interstitial maternal cytoplasm of the zoosporangium (Fig. 5F-G). The transcriptome of the mature zoosporangia stage showed an upregulation of phagosome genes relative to the free-swimming zoospore stage (Fig. 5I). The zoospore vacuoles contained electron-dense cargo similar to lipids (Fig. 5F). The prominence of this engulfment across replicates suggests that endocytosis is the primary mode by which resources are trafficked from the maternal cytoplasm into developing zoospores post-cleavage, and that zoospore development does not cease once cleavage has been completed. Notably, developing zoospores did not yet display a detectable ribosomal cluster, as in the free-swimming zoospores (Fig. 2), and the only KEGG categories higher in free-swimming zoospores than in the mature zoosporangia samples were associated with ribosomes (Fig. 5J), indicating that this structure is formed later in zoospore development than captured here. The apparent importance of maintaining ribosomes in the biology of zoospores closes the chytrid life cycle when considered with our early discussion on the distinctiveness of zoospores in the zoospore-germling transition.

## DISCUSSION

This study into the cellular and molecular biology of *R. globosum* has generated a developmental atlas of an archetypal chytrid life cycle, shedding light on the cell patterns of major life stages and the biological processes governing the transitions between them. Our key findings are summarised in (Fig. 6).

**Figure 6.**
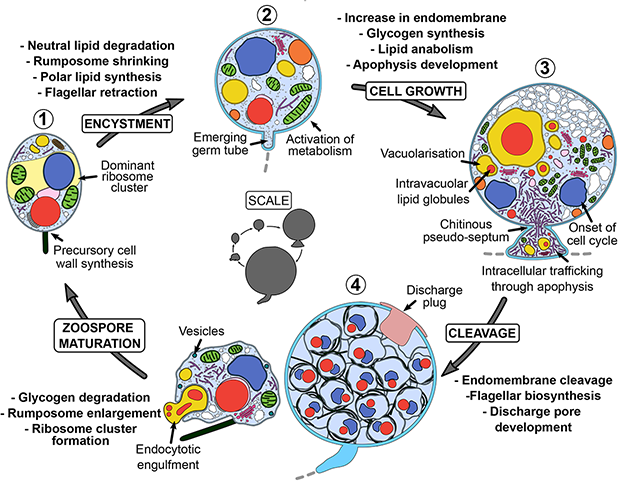
Summary of key components of the chytrid cell plan and biological processes associated with the transition between stages in the *R. globosum* life cycle. Inner life cycle shows life stages to scale. Grey dashed lines indicate the beginning of the rhizoid system.

In the *R. globosum* zoospore cell body, the ribosome cluster is a distinctive and dominating feature. Historically called the ‘nuclear cap’, ribosome clusters have been observed in zoospores throughout the Chytridiomycota (e.g. Koch, 1961) and in the closely related Blastocladiomycota (e.g. Lovett 1963). Lovett (1963) showed in *Blastocladiella* (Blastocladiomycota), as we show here with *R. globosum*, the ribosome cluster dissipates during the transition between the free-swimming zoospore and germling stages causing the release of the previously contained ribosomes throughout the cell. Lovett (1963, 1968) related the *Blastocladiella* zoospore ribosome cluster and subsequent dissipation with the biological activity of the cell during the zoospore-germling transition, proposing the role of the cluster was to maintain the ribosomes through the zoospore stage and to spatially isolate the ribosomes to prevent translation occurring until when the cluster dissipates (i.e. the germling stage) and protein synthesis is initiated. Other investigations into protein synthesis in chytrids (Léjohn and Lovett, 1965) and the Blastocladiomycota (Lovett, 1968; Schmoyer and Lovett, 1969) also suggest that translation does not begin until germination and that the zoospore is at least partially dependent on maternally-provisioned mRNA and ribosomes.

Our transcriptome data add molecular detail to this process, with KEGG categories related to ribosome maintenance downregulated and categories associated with translation and biosynthesis upregulated during the zoospore-germling transition. Similarly, (Rosenblum et al., 2008) detected high levels of transcripts associated with posttranslational protein modification in *Batrachochytrium dendrobatidis* (*Bd*) zoospores, but low transcriptional activity. Comparable translational activity is seen in dikaryan spore germination (Brambl and Van Etten, 1970; Mirkes, 1974; Rado and Cochrane, 1971). Taken together, the chytrid zoospore life stage represents a sophisticated and well-adapted specialised biological repertoire optimised for dispersal to new growth substrates or hosts rather than general metabolism, which is only initiated by the release of the ribosome cluster at the germling stage once favourable conditions are found.

The germling stage is characterised by major cell plan remodelling, including rhizoid growth, and concomitant activation of diverse metabolic pathways. Similar upregulation of metabolic pathways has been observed at the transcriptional level associated with conidial germination in dikaryan fungi (Sharma et al., 2016; Zhou et al., 2018). Interestingly, we detected the upregulation of proteasome genes in the germling relative to the zoospore, which are also necessary for dikaryan germination (Seong et al., 2008; Wang et al., 2011). A previous study into flagellar retraction in *R. globosum* showed that the internalised flagellum is disassembled and degraded in the germling stage, at least partially by proteasome-dependent proteolysis (Venard et al., 2020). Our findings of increased proteasome expression may likewise be associated with flagellar degradation and the recycling of redundant zoospore machinery in the germling.

The immature thallus displayed increased cellular and molecular signatures associated with the reproductive cell cycle, intracellular trafficking, and protein processing. A key structural development was the vacuolisation of the cell body. Highly vacuolated dikaryan cells (El Ghaouth et al., 1994; Gow and Gooday, 1987) are associated with diverse cellular processes including general homeostasis, protein sorting, cell cycling, and intracellular trafficking (Veses et al., 2008), any of which could at least partially explain the high vacuolisation of immature chytrid thalli. Noticeable in the context of chytrid cell biology however is the upregulation of actin-driven cytoskeletal genes, including those assigned to the Arp2/3 complex. The role of actin in vacuolisation and endocytosis has been demonstrated in yeast (Eitzen et al., 2002; Gachet and Hyams, 2005), similar to our observations here in *R. globosum*. Arp2/3-dependent actin dynamics drive crawling α-motility in some chytrid zoospores when moving freely in the environment (Fritz-Laylin et al., 2017; Medina et al., 2020) and the presence of animal-like actin components that have been lost in multicellular fungi makes chytrids useful models to investigate the evolution of the fungal cytoskeleton (Prostak et al., 2021). Although *R. globosum* zoospores do not crawl, the immature thallus has actin patches, cables, and perinuclear shells (Prostak et al., 2021). Here we show that, for a non-crawling chytrid, actin-associated genes are upregulated in immature thalli and are associated with a cell stage with high vacuolisation and endocytosis, which could possibly be associated with the early onset of zoosporogenesis.

We did not find any differences in molecular signatures related to cell wall synthesis between the different life stages at the higher categorical level in *R. globosum*, instead we observed individual differentially expressed genes suggesting that the process is dynamic and complex. Higher levels of putative chitin synthase gene transcripts (e.g. ORY39038) in wall-less zoospores was coupled with the detection of precursory cell wall material at the base of the flagellum. *Bd* transcriptomes also show specific transcripts associated with chitin synthesis to be higher in zoospores than in sessile thalli (Rosenblum et al., 2008). Chitin synthase activity has been shown associated with the *Blastocladiella emersonii* zoospore membrane (Dalley and Sonneborn, 1982). Early initiation of cell wall synthesis warrants further study and may explain why early chemical inhibition induces phenotypic disruptions to normal development in chytrids (Laundon et al., 2020). This emphasises the need to include the wall-less zoospore stage in investigations into chytrid cell wall biology.

This study has highlighted the complexity of lipid dynamics across the *R. globosum* lifecycle. Our data show that the volume of the lipid globule, total lipid by volume, and lipid as a percentage of dry mass remain unchanged between zoospores and germlings, yet we observed a shift in lipid type, moving from sphingolipids and neutral glycolipids (likely storage triacylglycerides) to polar lipids (likely membrane-associated phospholipids) between zoospores and germlings. Similarly, during the *B. emersonii* zoospore-germling transition glycolipids decrease and phospholipids increase (Dalley and Sonneborn, 1982). Previous research has characterised fatty acid profiles in chytrids (Akinwole et al., 2014; Gerphagnon et al., 2019; Rasconi et al., 2020) and shown differences between chytrid zoospores and sessile thalli of the same species (Taube et al., 2019). As the Nile Red emission spectrum undergoes a red shift in increasingly polar environments (Bertozzini et al., 2011), we propose that our live-cell data do not quantify the structural degradation of the lipid globule *per se* but rather biochemical polarisation as neutral storage lipids are catabolised and polar phospholipids are synthesised. The larger volumetric proportion of glycogen stores in developing zoospores over mature free-swimming zoospores also indicates that glycogen catabolism between the two stages contributes to the zoospore energy budget during motility as previously proposed (Powell, 1979).

Changes in lipid profiles were coupled with subcellular ultrastructure in *R. globosum*. The enzymatic function of LRM-associated microbodies as lipolytic organelles has been previously proposed (Powell, 1979, 1977, 1976), where evidence suggests that enzymatic activity increases following germination (Powell, 1976). From our data, this organelle may have bidirectional function and be associated with lipid production (anabolism and conversion) as well as catabolism. A key component of the LRM is the enigmatic chytrid rumposome, which was larger in zoospores than germlings. Previous hypotheses have proposed that this organelle is associated with environmental reception and signal transduction in flagellar regulation (Dorward and Powell, 1983). An enlarged rumposome in motile zoospores would support a flagellar role, but its retention in germlings implies additional functions, unless there is a prolonged delay in its degradation. The bulk of lipids in immature thalli during anabolism were intravacuolar and comparable intravacuolar inclusions have been identified in chytrid and dikaryan fungi in the past (Beakes et al., 1992; Bourett and Howard, 1994; Lösel, 1990). Intravacuolar lipid droplets have been previously investigated in yeast but in a catabolic capacity (Van Zutphen et al., 2014; Vevea et al., 2015). Although *de novo* storage lipid synthesis is associated with the ER (Vevea et al., 2015), the vacuoles identified here may cache and aggregate nascent globules as part of the lipid anabolic pathway.

The function of the chytrid apophysis has long been overlooked, despite its ubiquity in the Chytridiomycota (Powell, 1974; Powell and Gillette, 1987; Taylor and Fuller, 1980). Here we provide evidence that the apophysis is a distinct subcellular structure that acts as a junction for dynamic intracellular trafficking from the multiple branches of the rhizoid network into the central cell body. The localisation of high endomembrane activity in the apophysis and subsequent passage through the annular pore in the pseudo-septum into the cell body implicates this structure as a possible regulatory intermediary consolidating the rhizoid network. The ability of multicellular dikaryan fungi to translocate assimilated nutrients through their hyphal network from the site of uptake is sophisticated (van’t Padje et al., 2021; Whiteside et al., 2019), and the observed analogous endomembrane flow from feeding rhizoids to the cell body in chytrids is perhaps not surprising. However, the localisation of high endomembrane activity to the apophysis and through the pseudo-septum into the cell body implicates this structure as a regulatory and intermediary junction.

The pseudo-septation of the apophysis and rhizoids from the cell body is evidence for functional compartmentation (i.e. feeding vs. reproduction) within the thallus of a unicellular fungus. Comparable structures are also present in other chytrid species (e.g. Barr, 2011; Beakes et al., 1992). Division of multicellular dikaryan fungi by septa, where continuity between distinct cytoplasmic compartments is maintained by septal pores, is integral to multicellularity, cellular differentiation, and resilience (Bleichrodt et al., 2015, 2012). The origin of hyphal septa was a major innovation in fungal evolution (Berbee et al., 2017; Nagy et al., 2020) occurring at the node shared by hyphal and rhizoidal fungi (Berbee et al., 2017). The role of the apophysis/cell body pseudo-septum (or an analogous structure) in chytrids in delineating functionally dedicated subcellular compartments may represent an evolutionary precursor to dikaryan septa and differentiation. Therefore, investigating the chytrid apophysis is not only important for understanding intracellular trafficking biology in the phylum, but also the evolution of multicellularity more widely across the fungal kingdom.

Our quantitative reconstructions of individual developing zoospores in the zoosporangium and comparison with their free-swimming counterparts have added to understanding the underpinning biology of chytrid zoosporogenesis. Perhaps our most striking finding is the amoeboid morphology of developing zoospores, resulting from engulfment, and trafficking structures, suggesting that developing zoospores assimilate material from the maternal cytoplasm post-cleavage. Although dikaryan sporogenesis is complex and diverse, it typically involves the septation of hyphal cytoplasm via cell wall synthesis (Cole, 1986; Money, 2016). As walled cells, dikaryan spores are incapable of such engulfment activity and therefore amoeboid zoospores have more in common with their more distant opisthokont relatives in this regard. Nucleariid amoebae (Yoshida et al., 2009), choanoflagellates (Laundon et al., 2019), and various animal cell types (Bayne, 1990) exhibit analogous endocytotic engulfment behaviour as we show here in fungal zoospores. This apparent conservation may indicate that such engulfment behaviour to assimilate subcellular cargo during sporogenesis of wall-less zoospores existed in the last common ancestor of branching fungi and was lost in dikaryan fungi as spores became walled.

In conclusion, our characterisation of the *R. globosum* life cycle has revealed changes in cell structure and associated biological processes driving chytrid development, some of which show analogies in dikaryan fungi and others in ‘animal like’ cells. As important saprotrophs, parasites, and pathogens, our findings provide information into the cellular processes that underpin the ecological importance of chytrids. In addition, our characterisation of an early-diverging fungus that retains cellular characteristics from the last common ancestor of branching fungi is a step forward in reconstructing the putative biology of this organism. This study demonstrates the utility of developmental studies with model chytrids such as *R. globosum* and reiterates the need for fundamental biology in investigating the function of chytrid cells.

## METHODS AND MATERIALS

### Culture maintenance

*Rhizoclosmatium globosum* JEL800 was maintained on peptonised milk, tryptone, and glucose (PmTG) agar plates (Barr, 1986) in the dark at 23 ^°^C. To collect zoospores, mature plates were flooded with 5 ml PmTG and the cell suspension was passed through a 10 μm cell sieve (pluriSelect) to remove non-zoospore life stages. Zoospore density was quantified under a Leica DM1000 (10 x objective) with a Sedgewick Raft Counter (Pyser SCGI) diluted to 1:1000 and fixed in 0.2% formaldehyde. Zoospores were diluted to a working density of 3 x 10^6^ ml^-1^ prior to inoculation for all light microscopy experiments.

### Cell harvesting for SBF-SEM, transcriptomics, and lipid quantification

*R. globosum* was grown to progress through the life cycle and sampled at key time points: 0 h (zoospore), 1.5 h (germling), 10 h (immature thalli), and at 24 h when the population was a mix of stages including mature zoosporangia (Fig. 1C, Suppl. Fig. 8). For zoospores, each replicate was harvested from 10 ml of undiluted cell suspension immediately after plate flooding. For germlings, each culture flask (83.3910, Sarstedt) contained 40 ml of liquid PmTG and was inoculated with 10 ml of zoospore suspension, incubated for 1.5 h, and pelleted after scraping the flask with an inoculation loop to dislodge adherent cells. Immature thalli replicates were pooled from 10 x culture flasks of 25 ml liquid PmTG inoculated with 50 µl zoospore suspension and incubated for 10 h. Mixed 24 h populations containing mature zoosporangia were harvested and strained through a 40 µm cell strainer (11587522, FisherBrand) to remove smaller life stages. All incubations were conducted at 23 °C and cells were pelleted at 4,700 rpm for 5 min. For SBF-SEM, cell pellets were resuspended and fixed in 2.5% glutaraldehyde in 0.1 M cacodylate buffer pH 7.2. Cells were harvested identically for RNA Seq (*n* = 3) with the exception that the supernatant was removed before being flash frozen in liquid nitrogen and stored at −80 °C. Sub-samples from cell pellets were diluted 1:1000, fixed in 0.2% formaldehyde, and stained with FM 1-43FX to visualise cell membranes in order to qualitatively confirm the synchronicity of cultures under a confocal microscope (see further below) before being processed further (Suppl. Fig. 8). Cell pellets were harvested for lipid extraction and quantification as per RNA samples (*n* = 3).

### SBF-SEM imaging and reconstruction

Samples were further fixed in buffered glutaraldehyde, pelleted, and embedded in either agar or Bovine Serum Albumin (BSA) gel. Blocks were processed into resin using a modified protocol by Deerinck and colleagues (https://tinyurl.com/ybdtwedm). Briefly, gel-embedded chytrids were fixed with reduced osmium tetroxide, thiocarbohydrazide, and osmium tetroxide, before being stained with uranyl acetate and lead aspartate. Stained blocks were dehydrated in an ethanol series, embedded in Durcupan resin, and polymerised at 60 °C for 24-48 h. Blocks were preliminarily sectioned to ascertain regions of interest (ROIs) using transmission electron microscopy (FEI Tecnai T12 TEM). ROIs were removed from the resin blocks and remounted on aluminium pins, which were aligned using scanning electron microscopy (Zeiss GeminiSEM) on a Gatan 3 view serial block face microtome and imaged.

Stacks of chytrid cells were acquired at 75 nm z-intervals with an XY pixel resolution of 2 nm (zoospore, germling, and developing zoospore inside a mature zoosporangium), 4 nm (immature thallus), and 8 nm (mature zoosporangium). Although XY pixel size differed between life stages, 2-4 nm resolutions were above the minimum sampling limits for quantitative comparison of reconstructed organelles. Due to the lack of replication, the mature zoosporangium was only considered qualitatively. Acquired stacks were cropped into individual cells (*n* = 5) and imported into Microscopy Image Browser (MIB) (Belevich et al., 2016) for reconstruction. Prior to segmentation, images were converted to 8-bit, aligned, contrast normalised across z-intervals using default parameters, and then were processed with a Gaussian blur filter (sigma = 0.6). Stacks were segmented using a combination of manual brush annotation and the semi-automated tools available in MIB (Suppl. Fig. 1). Briefly, flagella, lipids, microbodies, nuclei, ribosomal clusters, rumposomes, peripheral bodies, striated inclusions, vacuoles, and vesicles were segmented manually using interpolation every 3-5 slices where appropriate; the discharge plug, endomembrane, glycogen granules, Golgi apparatuses, and mitochondria were masked by coarse manual brushing and then refined by black-white thresholding; and cell boundaries were segmented using the magic wand tool. All models were refined by erosion/dilation operations and manually curated. Models were also refined by statistical thresholding at size cut-offs for each structure consistent across all life stages (either 500 or 1000 voxels).

Structures were volumetrically quantified within MIB. For visualisation of reconstructed cells .am model files were resampled by 33% in XY and imported as arealists into the Fiji (Schindelin et al., 2012) plugin TrakEM2 (Cardona et al., 2012), smoothed consistently across life stages, and exported as 3D .obj meshes for final rendering in Blender v2.79. All quantification was conducted on unsmoothed models scaled by 50%. Flagella and rhizoids were excluded from quantification as they are not a component of the cell body, and their total length were not imaged in this study. The unassigned cytosol fraction was defined as the total volume of assigned organelles subtracted from the total cell volume and is inclusive of small structures such as ribosomes, vesicles, and small endomembrane and glycogen objects that could not be confidently assigned and were conservatively excluded. Only endomembrane not considered to be predominantly structural (i.e. an organelle or cell-compartment boundary) was reconstructed in the endomembrane category.

### RNA extraction

RNA was extracted from the cell pellets using the RNeasy extraction kit (Qiagen) following the manufacturer’s instructions with minor modifications. Cell pellets were thawed in 600 ml RLT lysis buffer containing 10 μl ml^-1^ of 2-mercaptoethanol and lysed at room temperature for 5 min with periodic vortexing. Cell debris was removed by centrifuging at 8000 xg for 1 min, before the lysate was recovered and passed through a QIA shredder (Qiagen). An equal volume of 100 % ethanol was added to the homogenised lysate before being transferred to a RNeasy extraction column. RNA was then extracted following the manufacturers protocol and included an on-column DNase digestion step using the RNase-Free DNase (Qiagen). RNA was quantified using both a NanoDrop 1000 spectrophotometer (Thermo) and the RNA BR assay kit (Invitrogen) on the Qubit 4 fluorometer (Invitrogen). RNA quality was assessed using the RNA 6000 Nano kit total RNA assay (Agilent) run on the 2100 Bioanalyzer instrument (Agilent).

### Sequencing and bioinformatics

Sequencing was carried out using Illumina NovaSeq 6000 technology and base calling by CASAVA, yielding 20,122,633 – 23,677,987 raw reads by Novogene (www.novogene.com). Raw reads were filtered for adaptor contamination and low-quality reads (ambiguous nucleotides > 10% of the read, base quality < 5 for more than 50% of the read) resulting in 19,665,560 – 22,917,489 clean reads. Reads were mapped against the JEL800 genome using HISAT2 before differentially expressed genes (DEGs) between life stages were determined using DESeq2 as part of the Novogene pipeline (Love et al., 2014). Transcriptomic profiles were highly conserved between replicates within each of the three life stages (Suppl. Fig. 9). All further analyses were performed in house in R v3.6.1 (R Core Team) using output from the Novogene analysis pipeline. Shared genes between life history stages were displayed using UpSetR (Conway et al., 2017). Volcano plots of differentially expressed genes were produced using ggplot2 based upon a conservative threshold of log2FoldChange > 0, padj < 0.05. Gene Ontology (GO) and Kyoto Encyclopedia of Genes and Genomes (KEGG) enrichment analysis was carried out using the enricher function in the R package clusterProfiler v3.12 (Yu et al., 2012) with a threshold of padj < 0.05. Differentially expressed KEGG categories were plotted using the dotplot function (Fig. 2) and GO maps generated using the emapplot function (Suppl. Figs. 10-13). For the purposes of this study, analysis and discussion of KEGG pathways was favoured over GO categories as KEGG pathways allow for a more process-oriented interpretation of activity between the life stages.

### Confocal microscopy of subcellular structures

Cell structures were labelled in a 24 h mixed population with 5 µM calcofluor white (chitin), 1 µM Nile red (neutral lipid), and 5 µM FM1-43 (membranes). Cells were imaged under a 63 x oil immersion objective lens with a Leica SP8 confocal microscope (Leica, Germany). Image acquisition settings were as follows: for cell wall excitation at 405 nm and emission at 410-500 nm (intensity 0.1%, gain 20); for lipids excitation at 514 nm and emission at 550-710 nm (intensity 0.1%, gain 50); and for membranes excitation at 470 nm and 500-650 nm (intensity 5%, gain 50). All life stages were imaged under identical acquisition settings. Cell wall and lipid images are maximum intensity projections at 0.3 µm z-intervals and membrane images are single optical sections.

### Live-cell widefield microscopy

Time-lapse imaging of the development of fluorescently labelled subcellular structures was optimised for LED intensity and dye loads that did not interfere with normal cellular development relative to a no-dye control (Suppl. Fig. 14). Population-level development was imaged using an epifluorescent Leica DMi8 microscope (Leica, Germany) with a 20 x objective lens, and single-cell development with a 63 x oil-immersion lens. Cell structures were labelled as above, with the exception of 1 µM FM1-43 (membrane). Image acquisition settings were as follows: for cell wall excitation at 395 nm and emission at 435-485 nm (intensity 10%, FIM 55%, exposure 350 ms); for lipids excitation at 575 nm and emission at 575-615 nm (intensity 10%, FIM 55%, exposure 1 s); for membranes excitation at 470 nm and 500-550 nm (intensity 10%, FIM 55%, exposure 2 s); and bright field (intensity 15, exposure 150 ms). Images were captured using a CMOS Camera (Prime 95B™, Photometrics). 500 μl of diluted zoospore suspension was applied to a glass bottom dish and cells were allowed to settle in the dark for 15 min. After this, the supernatant was removed and 3.5 ml of dye-containing PmTG was added to the dish and imaged immediately. To prevent thermal and hypoxic stress during the imaging period, the dish was placed into a P-Set 2000 CT stage (PeCon, Germany) where temperature was controlled at 22 °C by an F-25 MC water bath (Julabo, Germany), and the dish was covered by an optically clear film which permits gas exchange. Single images were taken at 15 min intervals for a total of 18 h for population-level development, and 50 µm z-stacks (2 µm z-intervals) at an interval of 10 min for a total of 14 h for single-cell development. Lipid degradation in live, settled zoospores was likewise imaged using 100 µl zoospore suspensions labelled with Nile Red in glass bottom dishes. For comparison, labelled zoospores fixed in 0.2% formaldehyde were also imaged to control for photobleaching. Cells were imaged at 30 s intervals for 2 h. To visualise endomembrane trafficking in the apophysis, 100 µl of 24 h mixed PmTG cultures stained with 10 µM FM 1-43 were likewise imaged in glass bottom dishes under a 63 x oil immersion objective at 30 s intervals for 30 min.

### Image analysis for live-cell microscopy

Developmental time series of fluorescently labelled subcellular structures were analysed with a custom workflow (Suppl. File 3-4) based around scikit-image 0.16.2 (Van Der Walt et al., 2014) run with Python 3.7.3 implemented in Jupyter Notebook 6.0.3. Briefly, cells were segmented using the bright-field channel by Sobel edge detection (Kanopoulos et al., 1988) and Otsu thresholding (Otsu, 1979). This mask was used to quantify normalised intensity in the fluorescence channel. For lipid tracking during single-cell development, images from the lipid channel were converted to maximum intensity projections and lipid globules were automatically detected using differences of Gaussian (DoG) detection in the Fiji plugin TrakMate (Tinevez et al., 2017). Tracking of the initial lipid globule was conducted using a simple LAP tracker.

### Lipid extraction and quantification

Lipids were extracted using the Bligh and Dyer method (Bligh and Dyer, 1959). Lyophilised culture pellets were submersed in a 2:1:0.8 (v/v/v) methanol (MeOH), dichloromethane (DCM) and phosphate-buffer (PB) and sonicated for 10 min in an ultrasonic bath before being centrifuged at 3,000 rpm for 2 min. The supernatant was collected, and the pellet was re-extracted twice. The combined supernatant was phase separated via addition of DCM and PB (giving an overall ratio of 1:1:0.9 (v/v/v)) and centrifugation at 3000 rpm for 2 min. The lower solvent phases were extracted prior to washing the remaining upper phase twice with DCM. The three lower solvent phases were collected and gently evaporated under oxygen-free nitrogen (OFN) in a water bath held at 25 °C (N-EVAP, Organomation, USA). The initial lipid extracts were weighed to quantify total lipid biomass before being dissolved in 9:1 (v/v) DCM:MeOH and loaded onto preactivated silica gel (4 h at 150 °C) columns for fractionation. Lipid fractions were separated by polarity via washing the column with one volume of DCM, followed by one volume of acetone and two volumes of MeOH. Each fraction was collected separately, evaporated to dryness under OFN and weighed.

### Statistical analysis

All data were tested for normality and homogeneity assumptions using a Shapiro and Levene’s test respectively. If assumptions could be met, then differences between zoospore, germling, and immature thallus volumetric proportions were assessed using ANOVA followed by Tukey HSD posthoc testing, or if not, then by a Kruskal-Wallis followed by a Dunn’s posthoc test. If a structure was entirely absent from a life stage (e.g. no cell wall in the zoospore stage) then the life stage was eliminated from statistical analysis to remove zero values and the remaining two life stages were compared using a *t*-test or Mann-Whitney U test depending on assumptions, and then the removed life stage qualitatively assigned as different. The differences between cell bodies and apophyses in the immature thallus life stage, and between mature zoospores and developing zoospores, were compared using either a paired *t*-test or a Mann-Whitney U test depending on assumptions. All statistical analysis was conducted using the scipy package (Virtanen et al., 2020) run with Python 3.7.3 implemented in Jupyter Notebook 6.0.3.

## ACKNOWLEDGEMENTS

We thank Chris Neal at the Wolfson Bioimaging Facility (University of Bristol) for SBF-SEM optimisation and imaging acquisition. We would also like to thank Joyce Longcore (University of Maine) for providing *R. globosum* JEL800 from her chytrid culture collection (now curated by the Collection of Zoosporic Eufungi at the University of Michigan https://czeum.herb.lsa.umich.edu).

## FUNDING

DL is supported by an EnvEast Doctoral Training Partnership (DTP) PhD studentship funded from the UKRI Natural Environment Research Council (NERC grant no. NE/L002582/1). NC, KE, ST and MC are supported by the European Research Council (ERC) (MYCO-CARB project; ERC grant agreement no. 772584).

**AUTHOR CONTRIBUTIONS**

DL and MC conceived and designed the study. DL setup the study, collected the samples and performed all microscope-based analysis (except the SBF-SEM optimisation and imaging acquisition). KB extracted the RNA and prepared for transcriptome sequencing. ST performed the lipid analysis. NC processed the transcriptome data and with TM supported associated data interpretation. DL and MC, with the help of NC, interpreted the data from the study. DL and MC wrote the manuscript with input from all the co-authors.

## DATA AVAILABILITY

All 3D objects, raw datasets associated with figures, and image analysis scripts can be found as Suppl. Files 1-4, in addition to all movies, processed SBF-SEM stacks, and model files, available for download from Figshare at: https://tinyurl.com/yww6h9d9.

Upon acceptance, raw sequencing reads will be deposited in the European Nucleotide Archive under ENA project PRJEB47366.

---

**Supplementary Figure 1.**
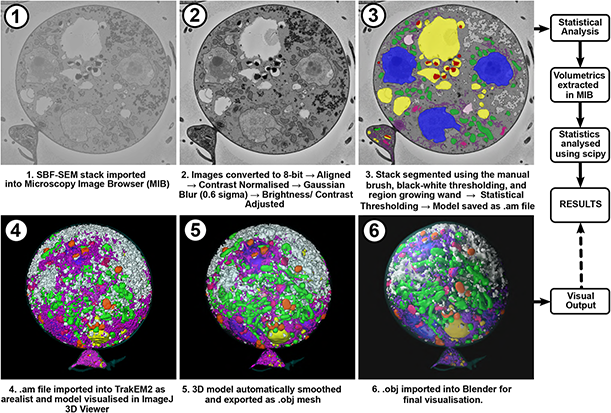
Workflow of the image analysis protocol used to generate and visualise 3D reconstructions of chytrid cells from SBF-SEM stacks.

**Supplementary Figure 2.**
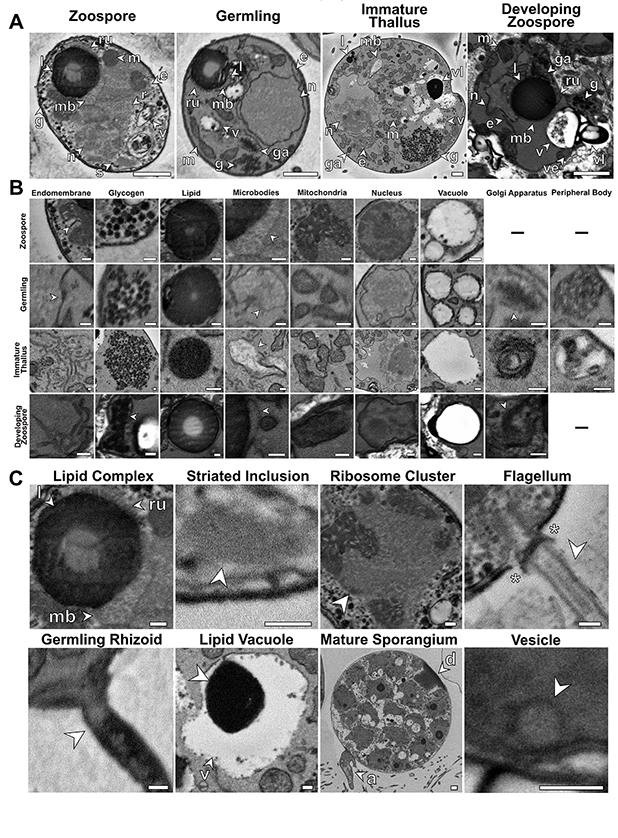
Examples of subcellular components identified in this study taken from single SBF-SEM slices. (A) Whole cell slices of each chytrid life stage showing the localisation and orientation of subcellular structures in context. (B) High magnification images of individual subcellular structures identified across life stages, where present. (C) Individual subcellular structures largely unique to individual life stages. a = apophysis, d = discharge plug, e = endomembrane, g = glycogen, ga = Golgi apparatus, l = lipid globule, m = mitochondria, mb = microbodies, n = nucleus, r = ribosomal cluster, ru = rumposome, s = striated inclusion, v = vacuoles. Asterisks in (C) show electron-dense plate at the base of the zoospore flagella. Scale bars = 1 µm (A) and 0.2 µm (B-C).

**Supplementary Figure 3.**
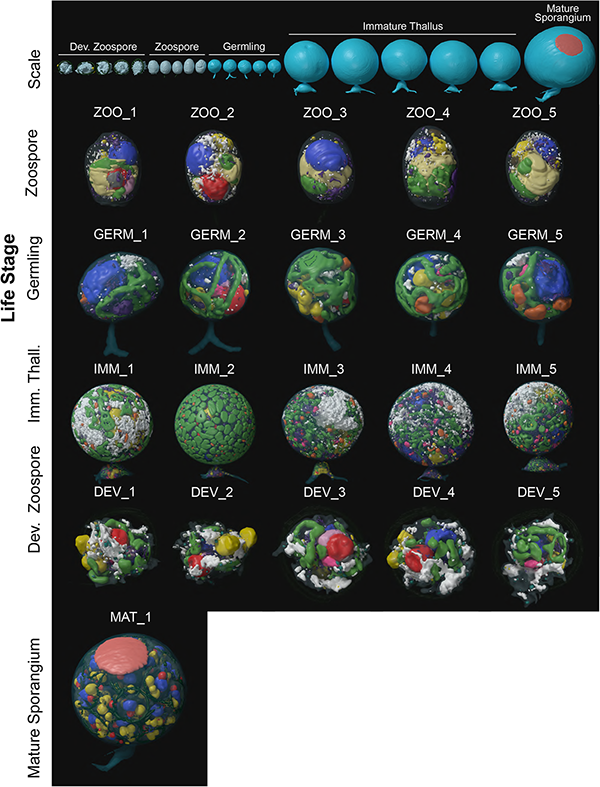
Individual 3D SBF-SEM reconstructions of *R. globosum* cells (*not to scale*) across life stages labelled with replicate ID’s. Organelle colours as in Fig. 2A-D. Top row shows all replicates to scale.

**Supplementary Figure 4.**
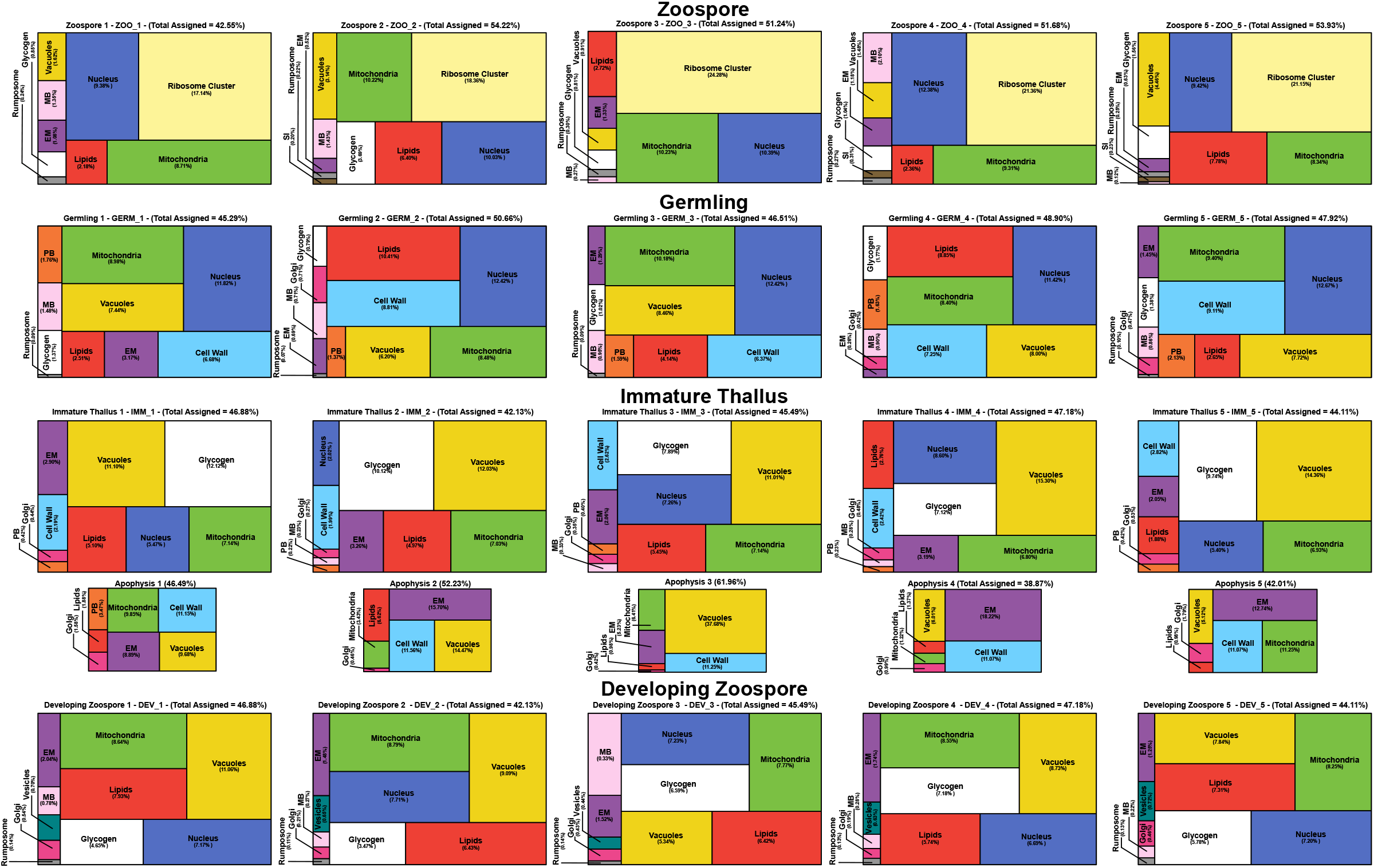
Individual volumetric compositions of assigned organelles from *R. globosum* SBF-SEM reconstructions across life stages labelled with replicate ID’s. Organelle colours as in Fig. 2A-D.

**Supplementary Figure 5.**
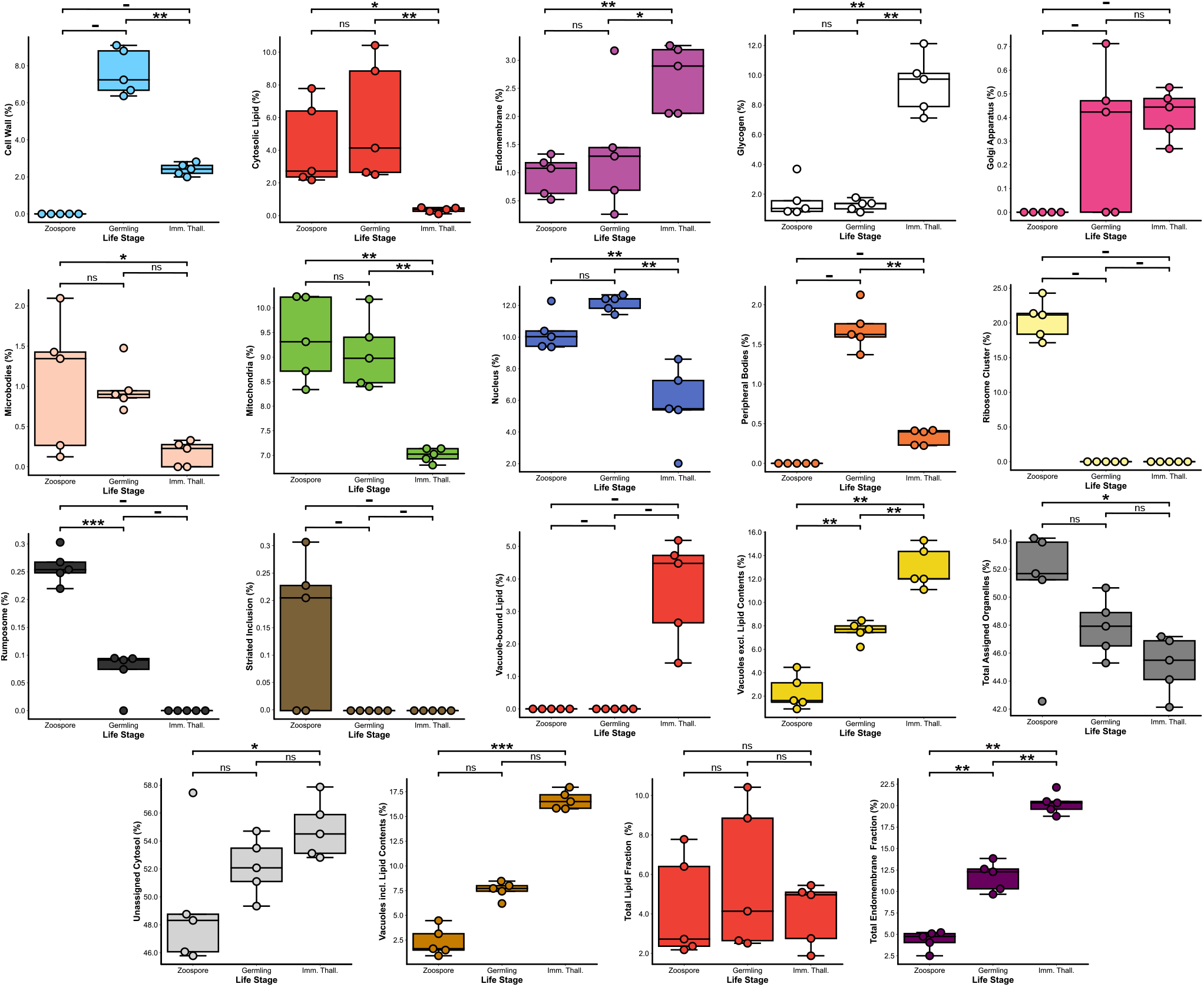
Comparisons of volumetric proportions of subcellular structures across chytrid life stages (*n* = 5). n.s *p* > 0.05 (not significant), \**p* < 0.05, \*\**p* < 0.01, \*\*\**p* < 0.001.

**Supplementary Figure 6.**
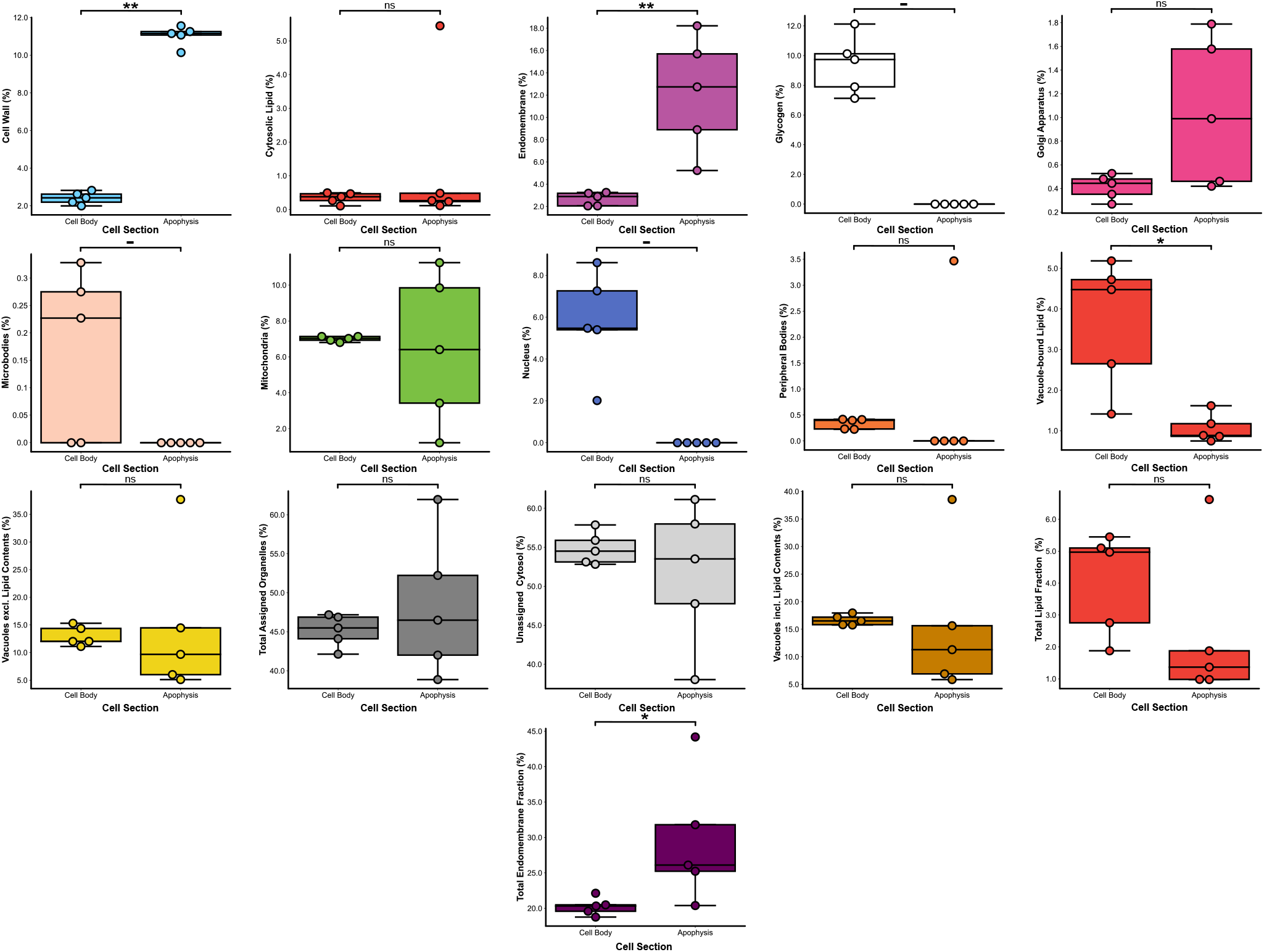
Comparisons of volumetric proportions of subcellular structures between immature thalli cell bodies and their corresponding apophyses (*n* = 5). n.s *p* > 0.05 (not significant), \**p* < 0.05, \*\**p* < 0.01, \*\*\**p* < 0.001.

**Supplementary Figure 7.**
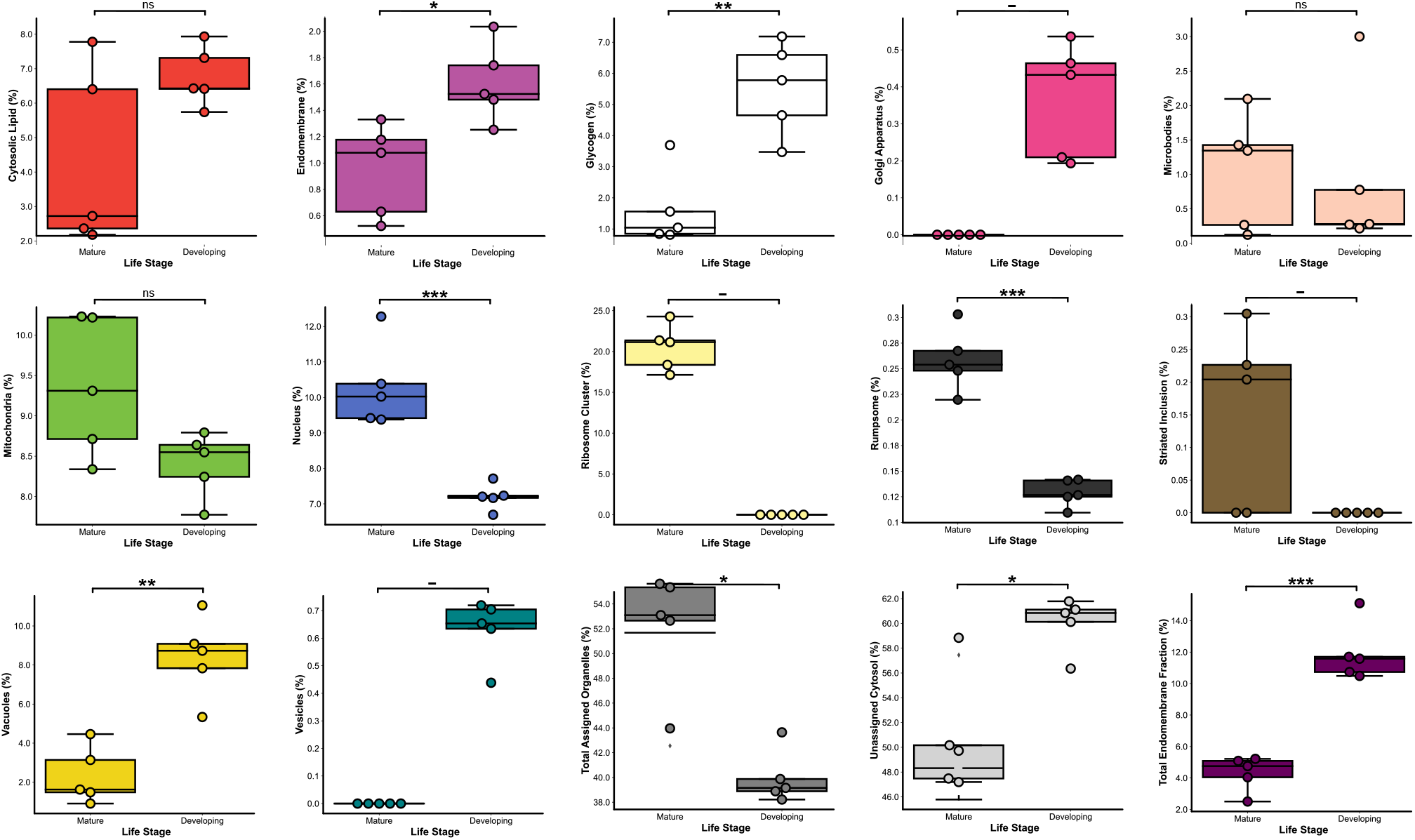
Comparisons of volumetric proportions of subcellular structures between developing and mature zoospores (*n* = 5). n.s *p* > 0.05 (not significant), \**p* < 0.05, \*\**p* < 0.01, \*\*\**p* < 0.001.

**Supplementary Figure 8.**
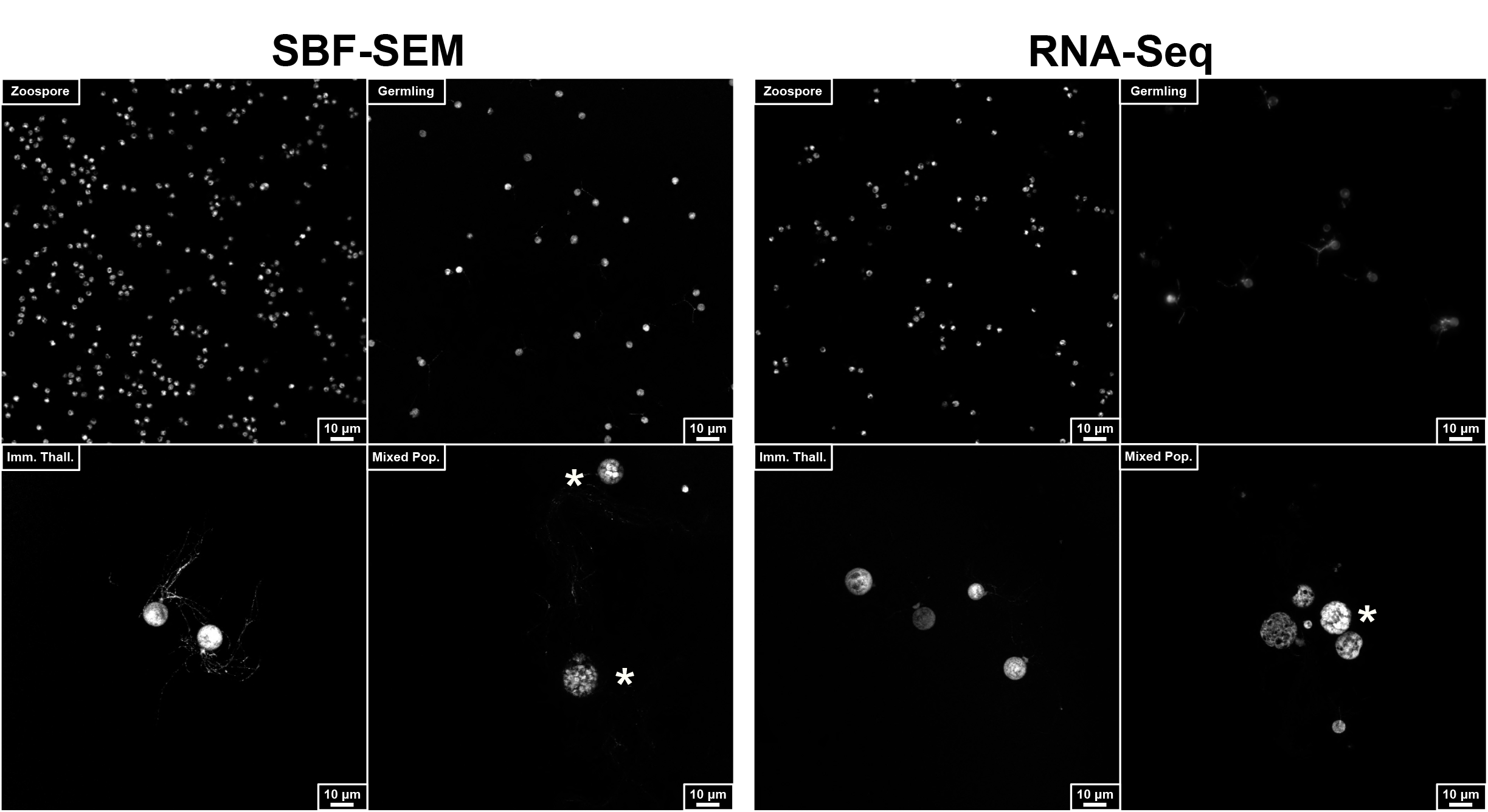
Representative images from confocal surveys conducted to assess the synchronicity of cell cultures for SBF-SEM and RNA-Seq harvesting. Cells diluted 1:1000, fixed in 0.2% formaldehyde, and stained with FM 1-43FX to visualise cell membranes. Asterisks mark mature zoosporangia in mixed populations.

**Supplementary Figure 9.**
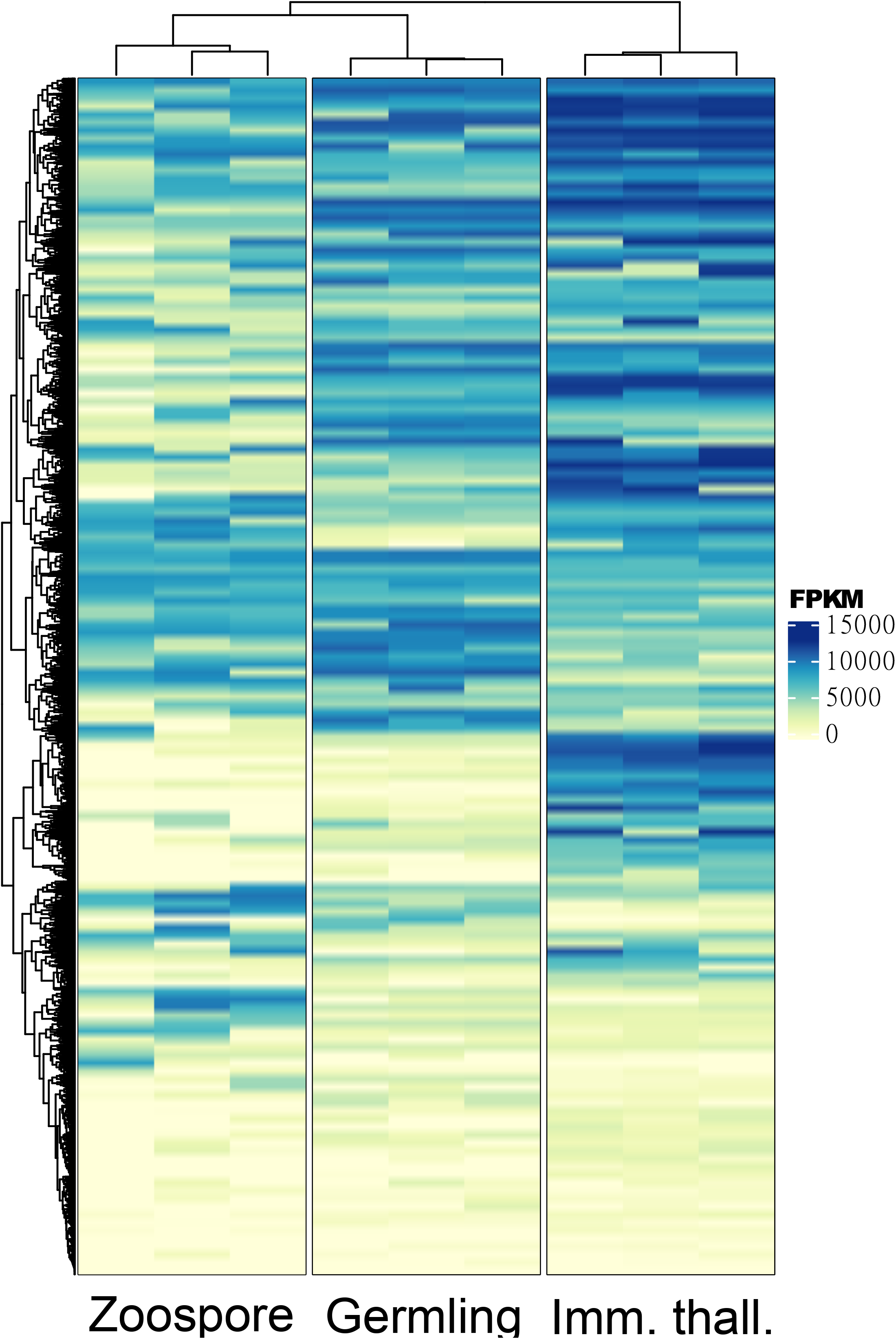
Heatmap clustering of all DEGs between zoospore, germling, and immature thallus replicates.

**Supplementary Figure 10.**
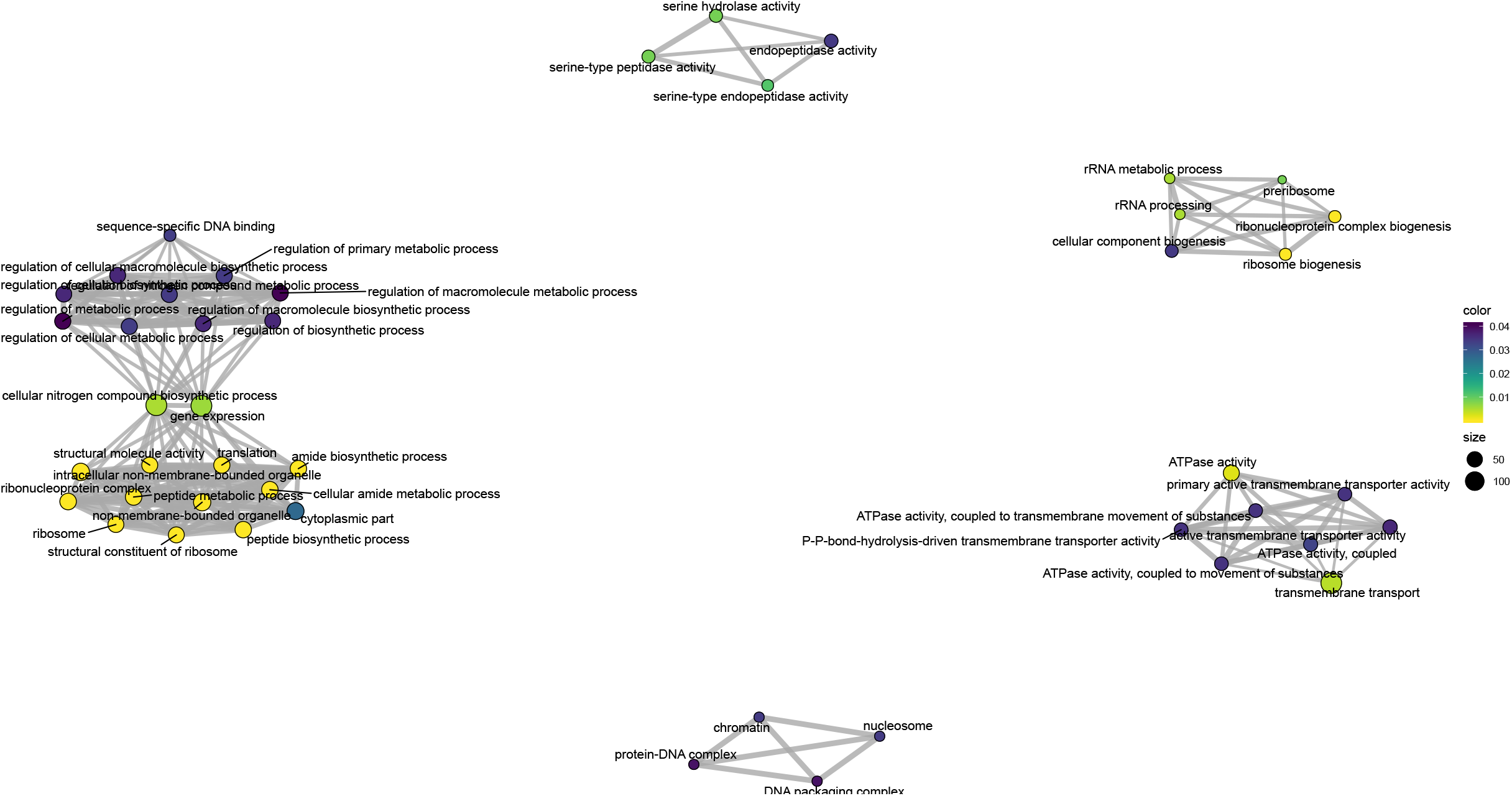
GO enrichment map showing significant (*p* < 0.05) differential expression of GO clusters, downregulated in germlings relative to zoospores. Circle size represents numbers of genes, colour represents adjusted *p*-value.

**Supplementary Figure 11.**
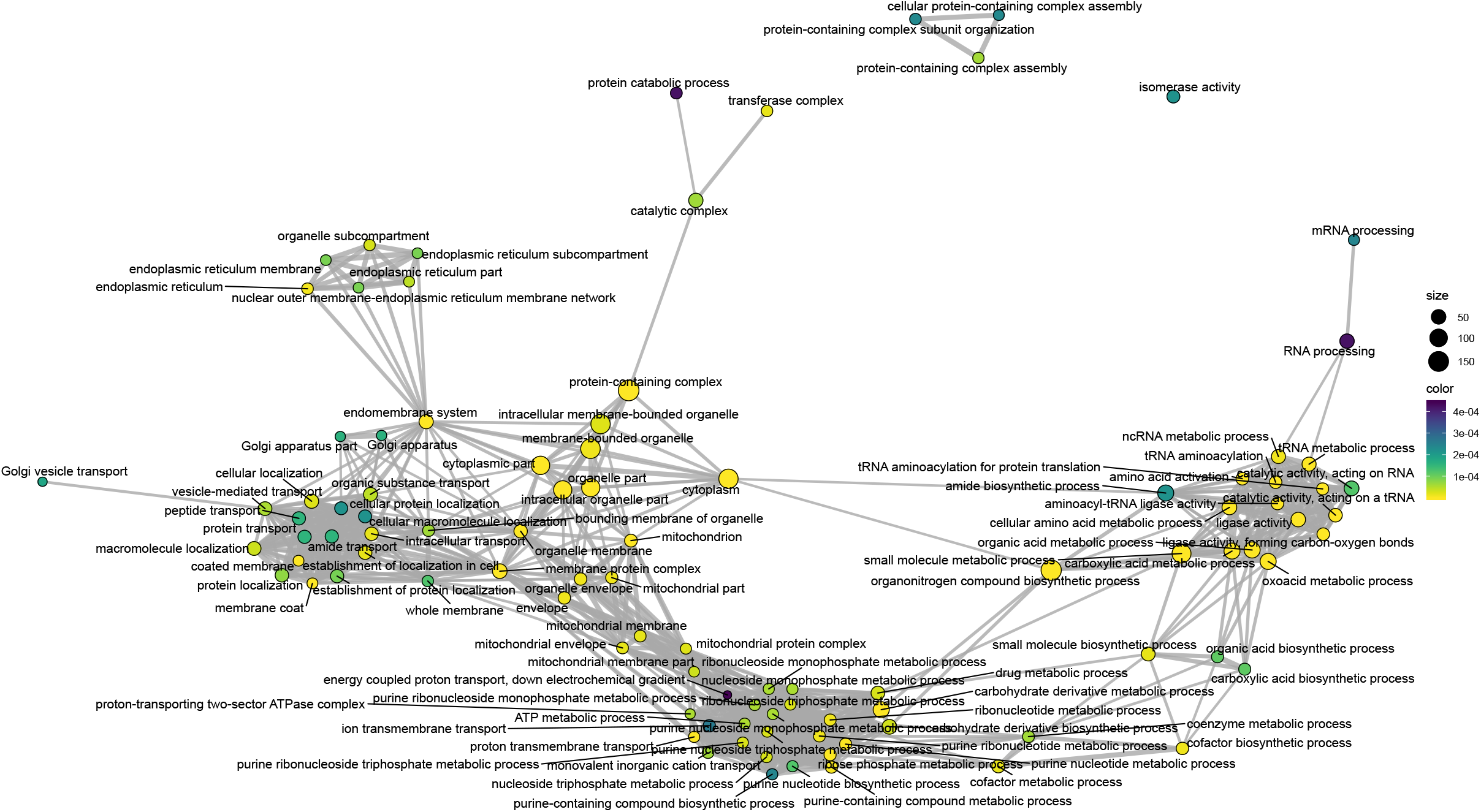
GO enrichment map showing significant (*p* < 0.05) differential expression of GO clusters, upregulated in germlings relative to zoospores. Circle size represents numbers of genes, colour represents adjusted *p*-value.

**Supplementary Figure 12.**
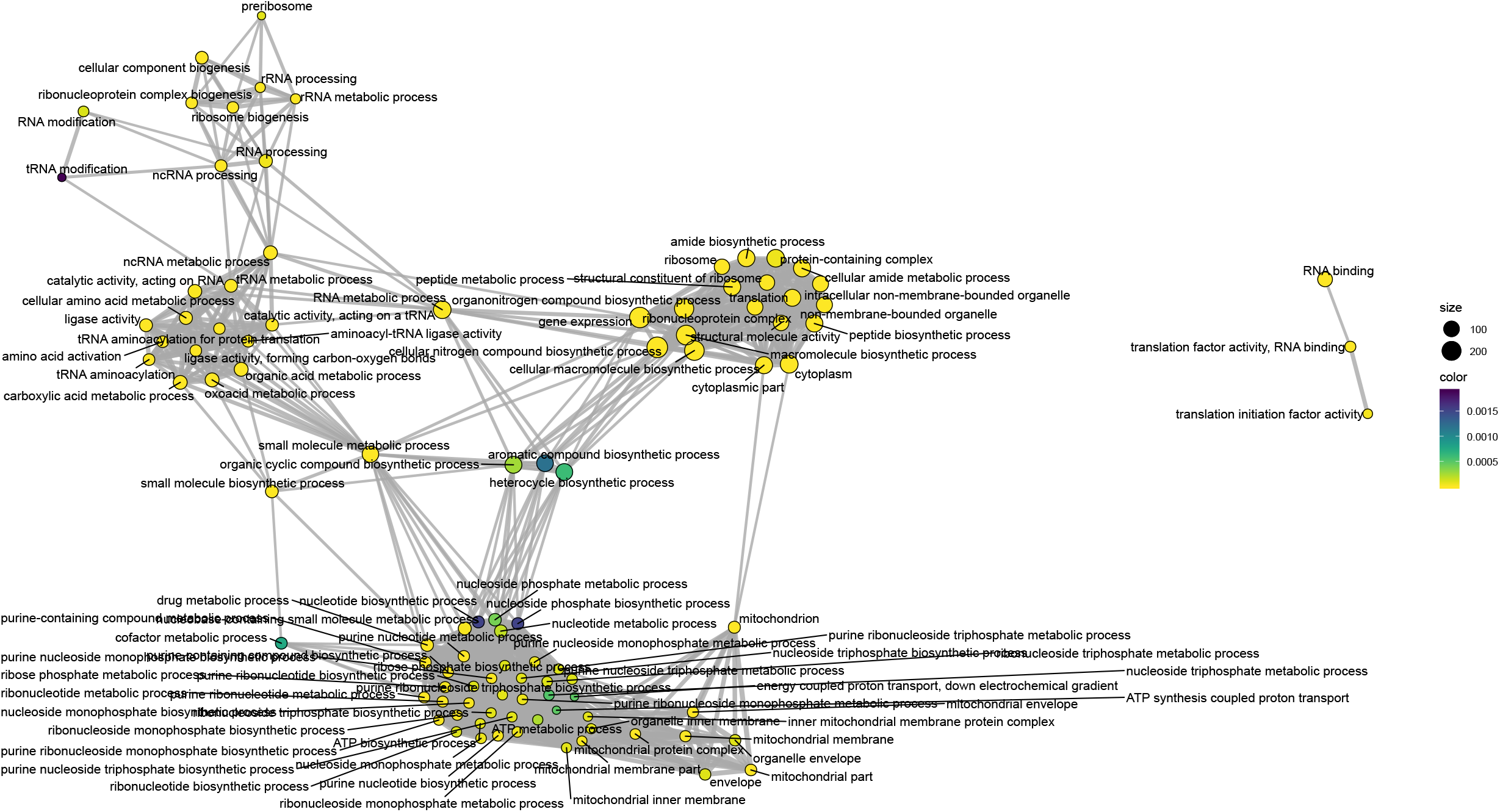
GO enrichment map showing significant (*p* < 0.05) differential expression of GO clusters, downregulated in immature thalli relative to germlings. Circle size represents numbers of genes, colour represents adjusted *p*-value.

**Supplementary Figure 13.**
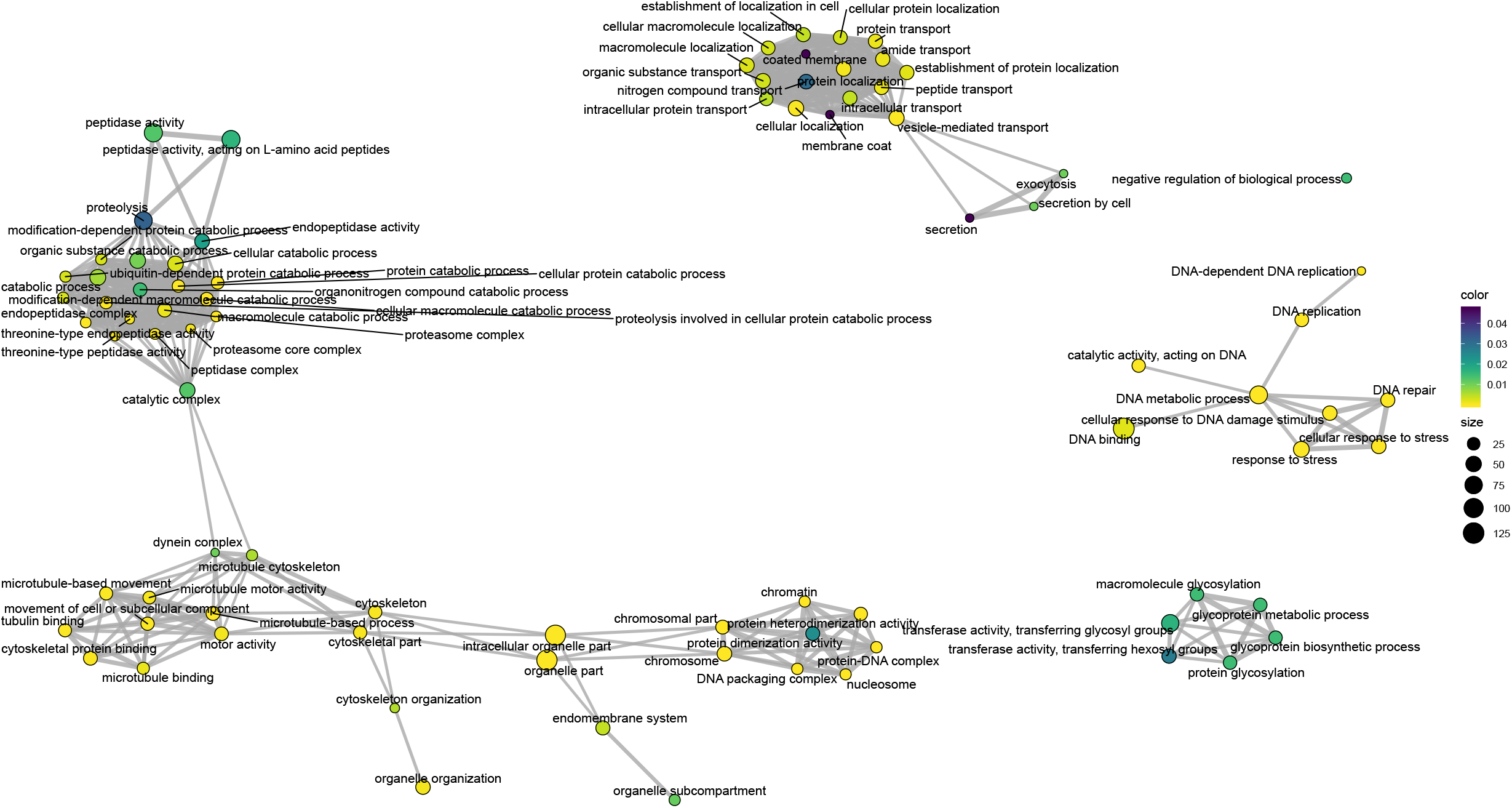
GO enrichment map showing significant (*p* < 0.05) differential expression of GO clusters, upregulated in immature thalli relative to germlings. Circle size represents numbers of genes, colour represents adjusted *p*-value.

**Supplementary Figure 14.**
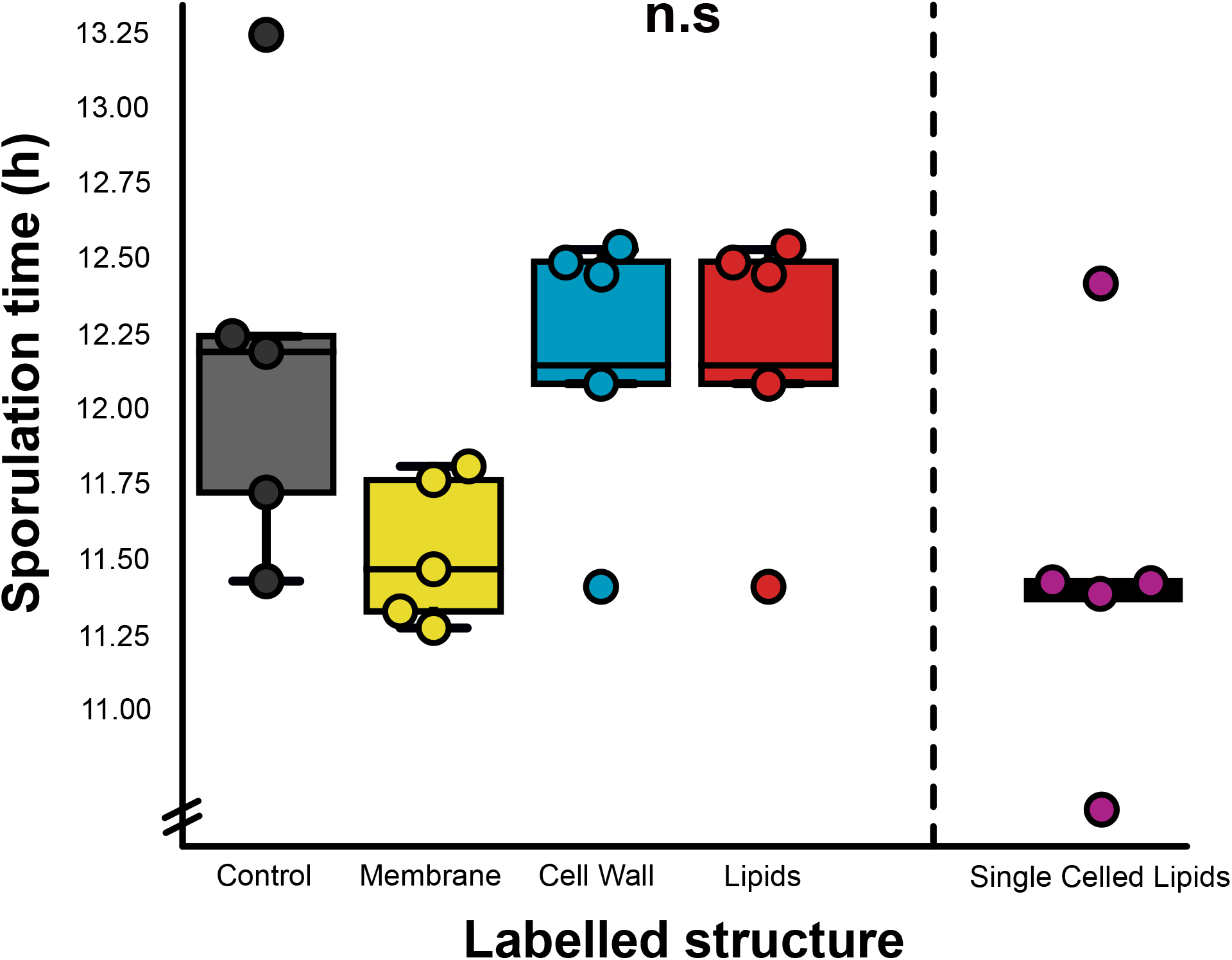
Comparison of sporulation times (as a proxy for normal cell development) for dye-labelled chytrid populations (*n* = 5) imaged by live-cell microscopy, relative to no dye-controls. n.s *p* > 0.05 (not significant).

## MOVIE LEGENDS

**Movie 1 SBF-SEM reconstructions allowed the structural comparison of life stages in *R. globosum.*** Representative SBF-SEM reconstructions of the zoospore, germling, and immature thallus life stages for comparison. Zoospore and germling cells shown to scale at the beginning of the movie, and later enlarged.

**Movie 2 Structural shifts in lipid globules were observed across *R. globosum* life stages, associated with the change from catabolism/conversion to anabolism.** Representative SBF-SEM reconstructions of the zoospore, germling, and immature thallus lipid structures for comparison.

**Movie 3 The zoospore lipid globule remained as an intact structure across the *R globosum* life cycle.** Automated particle tracking of lipid globules (red) across the chytrid lifecycle. Magenta circles mark individual lipid globules. Yellow track shows particle tracking of the initial lipid globule into the period of lipid anabolism. Cell wall shown in cyan. Timestamp = HH:MM.

**Movie 4 The *R.* globosum apophysis is structurally dominated by endomembrane structures.** Representative SBF-SEM reconstruction of a chytrid apophysis from an immature thallus.

**Movie 5 The *R.* globosum apophysis regulates intracellular trafficking between the rhizoids and cell body.** Live-cell imaging of endomembrane dynamics in the chytrid apophysis. The apophysis links endomembrane dynamics between the rhizoid system and thallus. Shown are DIC (left), endomembrane (centre), and overlay (right) channels. Timestamp = MM:SS.

**Movie 6 SBF-SEM reconstruction of an *R. globosum* mature zoosporangium.**

**Movie 7 Developing zoospores were more amoeboid than mature zoospores in *R.* globosum, due to elevated endocytosis and trafficking.** Representative SBF-SEM reconstructions of the ‘mature’ zoospore and developing zoospore life stages for comparison.

**Supplementary Movies 1-21** All individual SBF-SEM reconstructions used in this study. Replicates of zoospores (**Suppl. Mov. 1-5**), germlings (**Suppl. Mov. 6-10**), immature thalli (**Suppl. Mov. 11-15**), a mature zoosporangium (**Suppl. Mov. 16**), and developing zoospores (**Suppl. Mov. 17-21**).

**Supplementary Movie 22** Replicates of automated particle tracking of lipid globules (red) across the chytrid lifecycle. Magenta circles mark individual lipid globules. Yellow track shows particle tracking of the initial lipid globule into the period of lipid anabolism. Cell wall shown in cyan. Timestamp = HH:MM.

**Supplementary Movie 23** Replicates of live-cell imaging of endomembrane dynamics in the chytrid apophysis. The apophysis links endomembrane dynamics between the rhizoid system and thallus. Shown are DIC (left), endomembrane (centre), and overlay (right) channels. Timestamp = MM:SS.

## SUPPLEMENTARY TABLES

**Supplementary Table 1.** Volumetric quantities of cellular structures recorded across chytrid life stages.

| Cellular Structure | Chytrid Life stage – Volume in $\mu\text{m}^3$ | | | | | | | | | |
| --- | --- | --- | --- | --- | --- | --- | --- | --- | --- | --- |
|  | Zoospore<br>(n = 5) | ±<br>S.D | Germling<br>(n = 5) | ±<br>S.D | Immature<br>Thallus (n = 5) | ±<br>S.D | Imm. Thall.<br>Apophysis (n = 5) | ±<br>S.D | Dev. Zoospore<br>(n = 5) | ±<br>S.D |
| Total Volume | 20.749 | 1.687 | 33.991 | 2.042 | 1116.291 | 206.198 | 12.179 | 5.951 | 21.455 | 0.590 |
| Cell Wall | 0.000 | 0.000 | 2.594 | 0.425 | 26.448 | 3.112 | 1.326 | 0.600 | 0.000 | 0.000 |
| Cytosolic Lipid | 0.894 | 0.558 | 1.889 | 1.127 | 3.754 | 1.983 | 0.115 | 0.183 | 1.451 | 0.183 |
| Endomembrane | 0.195 | 0.070 | 0.479 | 0.410 | 30.829 | 11.485 | 1.390 | 0.729 | 0.346 | 0.072 |
| Glycogen | 0.332 | 0.253 | 0.433 | 0.138 | 104.359 | 27.516 | 0.000 | 0.000 | 1.190 | 0.334 |
| Golgi Apparatus | 0.000 | 0.000 | 0.106 | 0.101 | 4.498 | 0.898 | 0.153 | 0.149 | 0.079 | 0.033 |
| Microbodies | 0.226 | 0.189 | 0.336 | 0.118 | 2.025 | 1.857 | 0.000 | 0.000 | 0.194 | 0.253 |
| Mitochondria | 1.937 | 0.170 | 3.092 | 0.347 | 78.194 | 14.392 | 0.977 | 0.934 | 1.803 | 0.113 |
| Nucleus | 2.143 | 0.372 | 4.129 | 0.293 | 63.155 | 30.682 | 0.000 | 0.000 | 1.544 | 0.057 |
| Peripheral Bodies | 0.000 | 0.000 | 0.580 | 0.125 | 3.620 | 0.753 | 0.121 | 0.270 | 0.000 | 0.000 |
| Ribosome Cluster | 4.228 | 0.522 | 0.000 | 0.000 | 0.000 | 0.000 | 0.000 | 0.000 | 0.000 | 0.000 |
| Rumposome | 0.053 | 0.005 | 0.024 | 0.014 | 0.000 | 0.000 | 0.000 | 0.000 | 0.028 | 0.003 |
| Striated Inclusion | 0.032 | 0.031 | 0.000 | 0.000 | 0.000 | 0.000 | 0.000 | 0.000 | 0.000 | 0.000 |
| Vacuole-bound Lipid | 0.000 | 0.000 | 0.000 | 0.000 | 42.568 | 20.506 | 0.132 | 0.090 | 0.000 | 0.000 |
| Vacuoles excl. Lipid Contents | 0.488 | 0.311 | 2.576 | 0.369 | 144.077 | 31.251 | 1.568 | 1.234 | 1.809 | 0.472 |
| Vesicles | 0.000 | 0.000 | 0.000 | 0.000 | 0.000 | 0.000 | 0.000 | 0.000 | 0.135 | 0.024 |
| Total Assigned Organelles | 10.527 | 1.399 | 16.240 | 0.607 | 503.526 | 89.771 | 5.781 | 2.508 | 8.579 | 0.632 |
| Unassigned Cytosol | 10.222 | 1.268 | 17.751 | 1.670 | 612.764 | 122.699 | 6.398 | 3.675 | 12.877 | 0.389 |
| Vacuoles incl. Lipid Contents | 0.488 | 0.341 | 2.576 | 0.369 | 186.645 | 40.006 | 1.700 | 1.240 | 1.809 | 0.472 |
| Total Lipid Fraction * | 0.909 | 0.558 | 1.889 | 1.127 | 46.322 | 21.419 | 0.247 | 0.189 | 1.451 | 0.183 |
| Total Endomembrane Fraction ** | 0.275 | 0.375 | 4.018 | 0.802 | 227.616 | 51.755 | 3.363 | 1.212 | 2.563 | 0.445 |

*\*A functional category defined by the sum of cytosolic and vacuole-bound lipids.*
*\*\*A functional category defined by the sum of the endomembrane, Golgi apparatus, microbodies, peripheral bodies, vacuoles incl. lipid contents, and vesicles.*

**Supplementary Table 2.** Numerical quantities of cellular structures recorded across chytrid life stages.

| Cellular Structure | Chytrid Life stage – Volume in $\mu\text{m}^3$ | | | | | | | | | |
| --- | --- | --- | --- | --- | --- | --- | --- | --- | --- | --- |
|  | Zoospore<br>(n = 5) | ±<br>S.D | Germling<br>(n = 5) | ±<br>S.D | Immature<br>Thallus (n = 5) | ±<br>S.D | Imm. Thall.<br>Apophysis (n = 5) | ±<br>S.D | Dev. Zoospore<br>(n = 5) | ±<br>S.D |
| Total Volume | NA | NA | NA | NA | NA | NA | NA | NA | NA | NA |
| Cell Wall | 0.000 | 0.000 | 1.000 | 0.000 | 1.000 | 0.000 | 1.000 | 0.000 | 0.000 | 0.000 |
| Cytosolic Lipid | 1.000 | 0.000 | 1.000 | 0.000 | 68.800 | 55.233 | 13.600 | 14.276 | 1.000 | 0.000 |
| Endomembrane | 57.000 | 14.782 | 88.800 | 18.336 | 1513.800 | 641.545 | 45.000 | 35.050 | 88.400 | 21.629 |
| Glycogen | 331.200 | 76.424 | 167.800 | 56.193 | 1075.000 | 137.099 | 0.000 | 0.000 | 91.400 | 32.601 |
| Golgi Apparatus | 0.000 | 0.000 | 1.400 | 1.342 | 62.200 | 13.180 | 1.800 | 1.095 | 1.400 | 0.548 |
| Microbodies | 1.400 | 0.548 | 1.400 | 0.548 | 10.800 | 11.692 | 0.000 | 0.000 | 1.200 | 0.447 |
| Mitochondria | 2.800 | 2.490 | 1.200 | 0.447 | 237.200 | 129.820 | 27.600 | 31.198 | 9.000 | 3.082 |
| Nucleus | 1.000 | 0.000 | 1.000 | 0.000 | 1.800 | 1.304 | 0.000 | 0.000 | 1.000 | 0.000 |
| Peripheral Bodies | 0.000 | 0.000 | 3.600 | 2.074 | 52.600 | 23.650 | 1.200 | 2.683 | 0.000 | 0.000 |
| Ribosome Cluster | 1.000 | 0.000 | 0.000 | 0.000 | 0.000 | 0.000 | 0.000 | 0.000 | 0.000 | 0.000 |
| Rumposome | 1.000 | 0.000 | 0.8 | 0.447 | 0.000 | 0.000 | 0.000 | 0.000 | 1.000 | 0.000 |
| Striated Inclusion | 0.600 | 0.548 | 0.000 | 0.000 | 0.000 | 0.000 | 0.000 | 0.000 | 0.000 | 0.000 |
| Vacuole-bound Lipid | 0.000 | 0.000 | 0.000 | 0.000 | 70.600 | 39.835 | 4.400 | 2.074 | 0.000 | 0.000 |
| Vacuoles excl. Lipid Contents | 8.000 | 1.581 | 4.200 | 3.271 | 69.800 | 37.164 | 19.800 | 14.856 | 12.200 | 9.497 |
| Vesicles | 0.000 | 0.000 | 0.000 | 0.000 | 0.000 | 0.000 | 0.000 | 0.000 | 53.600 | 8.905 |
| Total Assigned Organelles | 405.000 | 81.557 | 272.200 | 59.302 | 3163.600 | 756.752 | 114.400 | 62.408 | 260.200 | 48.561 |
| Unassigned Cytosol | NA | NA | NA | NA | NA | NA | NA | NA | NA | NA |
| Vacuoles incl. Lipid Contents | 8.000 | 1.581 | 4.200 | 3.271 | 69.800 | 37.164 | 19.800 | 14.856 | 12.200 | 9.497 |
| Total Lipid Fraction * | 1.000 | 0.000 | 1.000 | 0.000 | 139.400 | 60.789 | 18.000 | 15.922 | 1.000 | 0.000 |
| Total Endomembrane Fraction ** | 75.400 | 19.424 | 99.400 | 19.501 | 1709.200 | 617.922 | 67.800 | 43.540 | 156.800 | 22.797 |

*\*A functional category defined by the sum of cytosolic and vacuole-bound lipids.*
*\*\*A functional category defined by the sum of the endomembrane, Golgi apparatus, microbodies, peripheral bodies, vacuoles incl. lipid contents, and vesicles.*

**Supplementary Table 3.** Volumetric percentages and statistical comparisons of cellular structures recorded across chytrid life stages.

| Cellular Structure | Chytrid Life stage – Volumetric % |  |  |  |  |  |  |  |  |
| --- | --- | --- | --- | --- | --- | --- | --- | --- | --- |
|  | Zoospore (n = 5) | ± S.D | Germling (n = 5) | ± S.D | Immature Thallus (n = 5) | ± S.D | Statistical Test used | p- Value | Posthoc Annotation |
| Total Volume | 100.000 | 0 | 100.000 | 0 | 100.000 | 0.000 | NA | NA | NA |
| Cell Wall | 0.000 | 0.000 | 7.644 | 1.245 | 2.409 | 0.328 | Mann Whitney U | <0.01 | A B C |
| Cytosolic Lipid | 4.290 | 2.610 | 5.714 | 3.678 | 0.341 | 0.159 | Kruskal | <0.01 | A A B |
| Endomembrane | 0.948 | 0.353 | 1.371 | 1.112 | 2.691 | 0.597 | ANOVA | <0.01 | A A B |
| Glycogen | 1.590 | 1.213 | 1.265 | 0.376 | 9.399 | 1.969 | Kruskal | <0.01 | A A B |
| Golgi Apparatus | 0.000 | 0.000 | 0.321 | 0.313 | 0.414 | 0.104 | Mann Whitney U | >0.05 | A B B |
| Microbodies | 1.052 | 0.836 | 0.978 | 0.293 | 0.167 | 0.156 | ANOVA | <0.05 | A AB B |
| Mitochondria | 9.363 | 0.861 | 9.086 | 0.732 | 7.005 | 0.143 | ANOVA | <0.001 | A A B |
| Nucleus | 10.297 | 1.187 | 12.151 | 0.512 | 5.749 | 2.477 | ANOVA | <0.001 | A A B |
| Peripheral Bodies | 0.000 | 0.000 | 1.696 | 0.278 | 0.336 | 0.100 | Mann Whitney U | <0.01 | A B C |
| Ribosome Cluster | 20.457 | 2.798 | 0.000 | 0.000 | 0.000 | 0.000 | NA | NA | A B B |
| Rumposome | 0.258 | 0.030 | 0.095 | 0.071 | 0.000 | 0.000 | T-Test | <0.001 | A B C |
| Striated Inclusion | 0.147 | 0.139 | 0.000 | 0.000 | 0.000 | 0.000 | NA | NA | A B B |
| Vacuole-bound Lipid | 0.000 | 0.000 | 0.000 | 0.000 | 3.689 | 1.596 | NA | NA | A A B |
| Vacuoles excl. Lipid Contents | 2.322 | 1.453 | 7.560 | 0.852 | 12.958 | 1.780 | ANOVA | <0.001 | A B C |
| Total Assigned Organelles | 50.724 | 4.754 | 47.857 | 2.082 | 45.159 | 2.087 | ANOVA | >0.05 | A AB B |
| Unassigned Cytosol | 49.276 | 4.754 | 52.143 | 2.082 | 54.841 | 2.087 | ANOVA | <0.05 | A AB B |
| Vacuoles incl. Lipid Contents | 2.322 | 1.453 | 7.560 | 0.852 | 16.647 | 0.930 | Kruskal | <0.01 | A A B |
| Total Lipid Fraction * | 4.290 | 2.610 | 5.714 | 3.678 | 4.030 | 1.604 | Kruskal | >0.05 | A A A |
| Total Endomembrane Fraction ** | 4.322 | 1.113 | 11.745 | 1.719 | 20.255 | 1.248 | ANOVA | <0.001 | A B C |

\*A functional category defined by the sum of cytosolic and vacuole-bound lipids.
\*\*A functional category defined by the sum of the endomembrane, Golgi apparatus, microbodies, peripheral bodies, vacuoles incl. lipid contents, and vesicles.

**Supplementary Table 4.** Volumetric percentages and statistical comparisons of cell bodies and their corresponding apophyses in immature thalli.

| Cellular Structure | Cellular Structure – Volumetric % |  |  |  |  |  |
| --- | --- | --- | --- | --- | --- | --- |
|  | Cell Body<br>(n = 5) | ±<br>S.D | Apophysis<br>(n = 5) | ±<br>S.D | Statistical Test used | p- Value |
| Total Volume | 100.000 | 0.000 | 100.000 | 0.000 | NA | NA |
| Cell Wall | 2.409 | 0.328 | 11.034 | 0.534 | Mann Whitney U | <0.01 |
| Cytosolic Lipid | 0.341 | 0.159 | 1.308 | 2.318 | Mann Whitney U | >0.05 |
| Endomembrane | 2.691 | 0.597 | 12.155 | 5.202 | Mann Whitney U | <0.01 |
| Glycogen | 9.399 | 1.969 | 0.000 | 0.000 | NA | NA |
| Golgi Apparatus | 0.414 | 0.104 | 1.047 | 0.627 | Mann Whitney U | >0.05 |
| Microbodies | 0.167 | 0.156 | 0.000 | 0.000 | NA | NA |
| Mitochondria | 7.005 | 0.143 | 6.429 | 4.215 | Mann Whitney U | >0.05 |
| Nucleus | 5.749 | 2.477 | 0.000 | 0.000 | NA | NA |
| Peripheral Bodies | 0.336 | 0.100 | 0.693 | 1.551 | Mann Whitney U | >0.05 |
| Vacuole-bound Lipid | 3.689 | 1.596 | 1.056 | 0.350 | Mann Whitney U | <0.05 |
| Vacuoles excl. Lipid Contents | 12.958 | 1.780 | 14.589 | 13.420 | Mann Whitney U | >0.05 |
| Total Assigned Organelles | 45.159 | 2.087 | 48.312 | 9.136 | Paired T-Test | >0.05 |
| Unassigned Cytosol | 54.841 | 2.087 | 51.688 | 9.136 | Paired T-Test | >0.05 |
| Vacuoles incl. Lipid Contents | 16.647 | 0.930 | 15.645 | 13.371 | Mann Whitney U | >0.05 |
| Total Lipid Fraction * | 4.030 | 1.604 | 2.364 | 2.407 | Paired T-Test | >0.05 |
| Total Endomembrane Fraction ** | 20.255 | 1.248 | 29.542 | 9.135 | Mann Whitney U | <0.05 |

*\*A functional category defined by the sum of cytosolic and vacuole-bound lipids.*
*\*\*A functional category defined by the sum of the endomembrane, Golgi apparatus, microbodies, peripheral bodies, vacuoles incl. lipid contents, and vesicles.*

**Supplementary Table 5.** Volumetric percentages and statistical comparisons of free-swimming and developing zoospores.

| Cellular Structure | Chytrid Life stage – Volumetric % |  |  |  |  |  |
| --- | --- | --- | --- | --- | --- | --- |
|  | Mature Zoospore<br>(n = 5) | ±<br>S.D | Developing Zoospore<br>(n = 5) | ±<br>S.D | Statistical Test used | p- Value |
| Total Volume | 100.000 | 0.00 | 100.000 | 100.000 | NA | NA |
| Cytosolic Lipid | 4.290 | 2.610 | 6.766 | 0.859 | Mann Whitney U | >0.05 |
| Endomembrane | 0.948 | 0.353 | 1.607 | 0.296 | T-Test | <0.05 |
| Glycogen | 1.590 | 1.213 | 5.536 | 1.494 | T-Test | <0.01 |
| Golgi Apparatus | 0.000 | 0.000 | 0.367 | 0.156 | NA | NA |
| Microbodies | 1.052 | 0.836 | 0.909 | 1.191 | Mann Whitney U | >0.05 |
| Mitochondria | 9.363 | 0.861 | 8.400 | 0.403 | T-Test | >0.05 |
| Nucleus | 10.297 | 1.187 | 7.202 | 0.361 | T-Test | <0.001 |
| Ribosome Cluster | 20.457 | 2.798 | 0.000 | 0.000 | NA | NA |
| Rumpsome | 0.258 | 0.030 | 0.129 | 0.013 | T-Test | <0.001 |
| Striated Inclusion | 0.147 | 0.139 | 0.000 | 0.000 | NA | NA |
| Vacuoles | 2.322 | 1.453 | 8.410 | 2.082 | T-Test | <0.001 |
| Vesicles | 0.000 | 0.000 | 0.630 | 0.113 | NA | NA |
| Total Assigned Organelles | 50.724 | 4.754 | 39.956 | 2.145 | Mann Whitney U | <0.05 |
| Unassigned Cytosol | 49.276 | 4.754 | 60.044 | 2.145 | Mann Whitney U | <0.05 |
| Total Endomembrane Fraction ** | 4.322 | 1.113 | 11.923 | 1.856 | T-Test | <0.001 |

**\*\*A functional category defined by the sum of the endomembrane, Golgi apparatus, microbodies, peripheral bodies, vacuoles incl. lipid contents, and vesicles.**

## SUPPLEMENTARY FILES

**Supplementary File 1** All SBF-SEM reconstructions available as 3D objects.

**Supplementary File 2** Raw data associated with figures and in-text discussions presented in this study.

**Supplementary File 3** Python script used to quantify population level fluorescence of developing chytrid cells (Fig. 3E, Fig. 4A).

**Supplementary File 4** Python script used to quantify single-cell Nile Red fluorescence of settled chytrid zoospores (Fig. 3F).

